# Integrative multi-omic and phenotypic analysis of open raceway pond production of *Monoraphidium minutum* 26B-AM reveals distinct stress signatures for scale-up and infection

**DOI:** 10.1101/2025.09.04.673846

**Authors:** Georgios Kepesidis, Jenna Schambach, Daniel Yang, Elise Wilbourn, Thomas Sheffield, Tyler Eckles, Olivia Watt, Matthew P. Hirakawa, Todd W. Lane, Raga Krishnakumar

## Abstract

**Background:** Green microalgae, such as *Monoraphidium minutum* 26B-AM, have garnered significant commercial interest due to their high biomass production and lipid yield, providing promising candidates for various bioprocessing applications. However, the economic viability of large-scale algal cultivation in open raceway ponds is limited by biocontamination and environmental stressors, necessitating deeper understanding of the molecular mechanisms that underpin resilience and productivity in these systems. We hypothesized that the molecular signature associated with the cellular responses of *M. minutum* to environmental stressors will reveal critical information for the timely prediction of resilience and productivity in algal cultures within open pond systems.

**Results:** To test this hypothesis, we conducted a longitudinal multi-omic study, integrating transcriptomics, proteomics, metabolomics, and phenomics, to monitor the acclimation, growth dynamics, and pathogen responses of algal cultures in two 1000 L raceway ponds, before and after the introduction of a pathogen as a stressor. We identified a number of molecular patterns that correlate with changes in the algal environment, and we can track these changes within the ponds per time. Furthermore, we identify scale-up and infection-specific molecular pathways through integrated multi-omics, showing that most patterns are unique to each studied stressor/transition.

**Conclusions:** Ultimately, this study demonstrates the utility of multi-omics observations at scale, revealing unique signatures and laying the groundwork for developing molecular detection techniques and predictive models that can improve the sustainability and efficiency of large-scale algae biomass production.

## 1. Background

Microalgae are a polyphyletic group of unicellular photosynthetic aquatic organisms that have attracted commercial interest since the middle of the past century because of their potential to advance biotechnology and biomanufacturing processes. Compared to land plants, their biomass yield is greater [1], and their high content of useful metabolites, such as lipids, has increased their presence in multiple industrial bioprocesses, including the production of food, pharmaceuticals, cosmetics, bioplastics, and biofuels. The high production costs, however, curtail most of these applications, especially for low-cost and high-volume products such as biofuels, making the use of algae currently unfeasible and unsustainable and restricting their potential [2] [3] [4].

A low-energy and low-cost method for large-scale production of algal biomass, given the financial restrictions for algal cultivation mentioned earlier, is the paddlewheel-based open raceway pond cultivation system [5] [6]. These widely used systems allow the cultures to grow in direct relation to the surrounding environment and the local weather conditions. Additionally, these ponds can be established on land that is not suitable for terrestrial agriculture, thereby utilizing a resource that would otherwise remain unused.

Despite their advantages, a number of barriers preclude the overall economic viability of bioprocesses within these systems. One crucial barrier is the susceptibility of the cultures to environmental “intruders”, which threaten both the quality of the final product and the overall survival of the pond cultures [7]. These events lead to pond crashes and a significant increase in operational costs, strongly establishing the need for their timely prediction.

In combination with the growth of undesired microbial organisms, open pond systems are susceptible to unpredictable changes in the weather [8], as are other site-specific environmental factors, such as sand particles or dust, which change the composition of the culture media [9]. All these factors, which are directly related to the immediate exposure of the cultures to their biotic and abiotic surroundings, can lead to unpredictable outcomes in terms of biomass productivity [10]. In addition, the growth and productivity are not always aligned with the observations made at smaller scales [11]. Understanding the effects that all these variables have on culture outcomes is critical to optimally predict growth in industrial settings. Consequently, optimal prediction can guide the design of resilient culture systems, ultimately reducing the cost of each operation. Identification of exposure- and threat-agnostic biomarkers of stress as expressed by the microalgae themselves would allow for detailed monitoring and remediation of open cultures in real-time and understanding what molecular pathways might be involved in resilience.

In the current industrial landscape, enhancing the resilience of expansive open pond systems and optimizing our predictive tools are essential for maintaining the stability of algal production. Here, we utilized cells as sensors of their environment by performing an in-depth multi-omics analysis (transcriptomics, proteomics, metabolomics and phenomics) in order to further understand the molecular dynamics behind the acclimation, growth, and response to infection by a parasitoid amoeboaphelid species (PL3) that are known to cause pond crashes, in two 1000 L open raceway ponds under environmentally-simulated conditions. For this study, we used the freshwater green microalga *Monoraphidium minutum* 26B-AM, a species currently studied as a promising winter species for biofuel feedstock because of its elevated lipid yield [12] [13] [14] [15] [16]. The genome of *M. minutum* has also been recently annotated [17]. [18, 19] Our work reveals a number of molecular signatures and regulatory elements of interest, and lays the groundwork for proactive interventions that can sustain algal health and performance in the face of environmental challenges. In this study, in addition to phenotypical observations, molecular markers evaluated via multi-omics are indicative of distinct stages of open pond culture. We verified this method by demonstrating a known link between light and redox signaling and photosynthesis and translation-related genes in *M. minutum*, in accordance with similar observations in closely related green algal species, confirming the applicability of the markers identified here across algal culture systems. Finally, we focused on the molecular pathways associated with the scale-up acclimation and the introduction of the pathogen into the system through integrated multi-omic analysis. While specific molecular networks of genes, proteins and metabolites were identified for each transition, we discovered unique signatures in response to the two stressors, revealing distinct signaling outcomes across stressors.

## 2. Methods

### 2.1 Strain and conditions

The *M. minutum* 26B-AM strain was initially collected and isolated as described previously [20], and it has been utilized in our laboratory as part of the DISCOVR strain library. Seed cultures of *M. minutum* (100 mL) were maintained in sterile 250 mL Erlenmeyer flasks, containing DISCOVR medium [20] under a 16:8 h light/dark cycle with continuous shaking at 110 rpm in a closed incubator with LED lighting. The light intensity during the light cycle was 100 µmol photons m^−2^ s^−1^ and the temperature was set between a maximum of 30 °C during the light period and a minimum of 23 °C during the dark period. The seed cultures were diluted regularly to maintain exponential phase growth.

### 2.2 Scale-up

The culture scale-up is graphically summarized in Fig.1. The seed cultures of *M. minutum* described above were used as inocula for 500 mL cultures grown in 1 L Erlenmeyer flasks. Progressively, the cultures were transitioned to 2 L bottles, 8 L carboys with stirring and air bubbling and 20 L carboys with bubbling. Each step lasted 10–15 days, until the inoculum was dense enough (approximately 2 million cells/mL) to be diluted to the next scale. Up to the 2 L scale, the cultures were grown under the same conditions as the initial seed cultures detailed in the previous section. For the 20 L carboys, the cultures were acclimated to relatively high light intensities (1500 µmol photons m^−2^ s^−1^). The 20 L carboys were thermally acclimated between 22-28 °C under a 12/12 h dark/light cycle in a water bath, to match the conditions of the next scale.

**Fig. 1:**
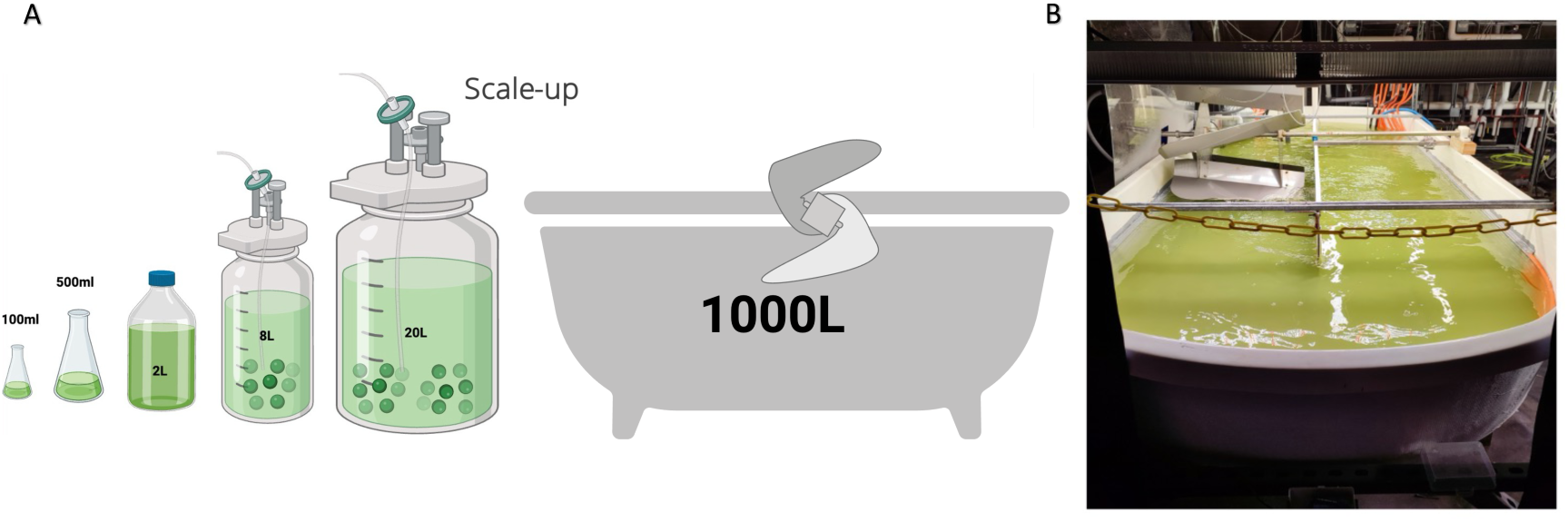
M.minutum scale-up. A) Graphic representation of the scale-up. B) Picture of the 1000L open raceway pond on Day 0 of the culture run, after the inoculation of the media with the precultures.

### 2.3 Cultivation and sample collection at the 1000 L opened raceway ponds

The cultures from the 20 L carboys (3 from each pond-total 60 L) were used as inocula for the 1000 L open raceway ponds. The ponds were filled with tap water and the nutrients and salts were added via the modified 10% BG-11 recipe, as described previously [20]. The ponds were operated under a 12 h/12 h dark/light cycle, implemented with LED light panels, placed over the ponds, reaching 1500 µmol photons m^−2^ s^−1^ during the light period, under a sinusoidal light intensity program (measured in closest proximity to the source). The temperature was maintained within a temperature range of 22-28 °C, with the highest temperature being reached in the middle of the light period and the lowest in the middle of the dark period. Temperature control was achieved through a heat-exchanger lining on the walls of the ponds. The cultures were constantly mixed with a mechanical paddlewheel at 20 rpm. The pH was held constant at 7.2 via CO_2_ sparging. The cells were grown under these conditions for 5 days and samples for the different omic analyses were collected on a daily basis one hour after the lights were turned on. For the phenotypic analysis, samples were collected across pond depths and locations to ensure that there were no extreme biases that were unaccounted for (more details below in 2.4). For the ‘omics datasets, samples were collected from the middle of the raceway ponds, at equal distance from the surface and the bottom of the pond and from the left and the right walls of the pond, under the assumption that this was the most successfully homogenous spot of the culture. The phenotypic and conditional measurements were taken directly after that.

On the 5^th^ day of the cultivation, 2/3 of each pond was drained and fresh water with new nutrients was added. In parallel, an inoculum of the parasitoid amoeboaphelid species FDO1 [19] [18] was added at a 0.05% volume concentration, in order to study the dynamics of *M.minutum* pond crashes caused by this strain. This specific strain has been isolated by the Arizona Center for Algae Technology and Innovation as the cause of *Monoraphdium minutum* pond crashes, and it has not yet been fully characterized. Under laboratory conditions, it caused *M. minutum* cultures to crash in 4-5 days. The determination of culture crashes was visual, when the ponds had completely transitioned from green to brown.

### 2.4 Phenotypic and conditional measurements

All phenotypic and conditional measurements were taken on a daily basis in technical triplicates unless otherwise stated. The cell count and cell size were determined with the Countess™ 3 FL Automated Cell Counter (Thermo Fisher Scientific, Waltham, MA, USA), from a sample collected from the ponds one hour after the lights were turned on (the maximal temperature and light intensity had been reached).

To measure chlorophyll fluorescence, 200 µL pond samples in 96-well transparent plates were measured at an excitation wavelength of 450 nm and an emission wavelength of 685 nm with a Spark® Multimode Microplate Reader (TECAN, Männedorf, Switzerland). The chlorophyll content of the ponds is presented in relative units.

For the ash-free dry weight (AFDW) measurements, the steps described in [21] were followed.

The total carbohydrate concentration was determined via the Total Carbohydrate Quantification Assay (Abcam, Cambridge, UK), following the manufacturer’s instructions. Samples were collected from each pond, alongside the samples for the omic analyses.

Samples were collected from each pond for the quantification of total H3 histones. This measurement was performed with the Histone H3 Total Quantification Kit (Abcam, Cambridge, UK), following the manufacturer’s protocol, and the colorimetric measurements were conducted using the Spark® Multimode Microplate Reader (TECAN, Männedorf, Switzerland).

Samples for the determination of caspase activity were also collected. For the estimation of cell lysis and caspase activity, the EnzChek Caspase-3 Assay Kit #1 (Thermo Fisher Scientific, Waltham, MA, USA) was used, with Z-DEVD-AMC as the substrate, according to the manufacturer’s protocol. Dissolved oxygen (%DO), pH, conductivity, temperature and light intensity were measured in each pond, immediately after sample collection. For each variable, we took the average of 9 collected data points, three from the surface, three from the middle and three from the bottom of the pond, in order to take into account the depth-based attenuation that is expected in some of those conditions. The %DO and temperature were measured with the Waterproof Dissolved Oxygen Meter Pen (Sper Scientific Direct, Scottsdale, AZ, USA), the pH and conductivity were measured with an Ohaus Starter AB33M1-F pH & Conductivity Bench Meter (Parsippany, NJ, USA), and the light intensity was measured with an LI-1500 Light Sensor Logger (Lincoln, NE, USA).

### 2.5 Community composition analysis

Samples were collected in sterile 15 mL Falcon tubes and stored at −20 °C before analysis. Prior to DNA extraction samples were thawed and media was removed by filtering through a filter plate (Cytiva AcroPrep, 3.0/0.2 µm pore size). Cells were resuspended with Zymo Bashing Bead Buffer and DNA was extracted following manufacturer instructions using a Zymo Quick-DNA Fungal/Bacterial 96 kit. Samples were amplified with the following primers: 357F and 783R [22] [23] for the 16S V3-V4 regions and TAReuk454 and TAReukREV3 TAReukREV3 [24] to amplify the 18S V4 region. Samples were indexed using Nextera XT indexes (Illumina, sets A and D) and 192 samples were multiplexed into a 4 nM final library. This library was sequenced using an Illumina MiSeq. Read classification and relative abundance calculations were performed in Python 3.13.5 using Kraken 2 and Bracken [25] [26] [27]. Community members were assigned using the Silva database (version 138.2) [28].

### 2.6 Transcriptomics/ RNA extraction and cDNA library preparation

The samples collected for the transcriptomic analyses (5 mL each) were centrifuged after collection at 5000 rpm for 2 minutes at 4 °C, and the resulting cell pellets were stored at −20 °C until RNA extraction was performed.

To initiate cell lysis, the frozen pellets were thawed on ice and resuspended in 750 µL of Qiagen RLT Buffer supplemented with 1% β-mercaptoethanol from the RNeasy Plant Mini Kit (Qiagen, Hilden, Germany). The resuspended cells were then transferred to 2 mL tubes containing Lysing Matrix Y zirconium oxide beads (MP Biomedicals, Santa Ana, CA, USA). Following a 10-second vortex, the samples were flash-frozen in liquid nitrogen. After thawing on ice, the cells were lysed via a FastPrep-24 bead beater (MP Biomedicals, Santa Ana, CA, USA) for 25 seconds at 40 Hz. The samples were subsequently centrifuged at 9000 rpm for 10 minutes at 4 °C, after which the clear supernatant was collected for RNA extraction. RNA was extracted via the RNeasy Plant Mini Kit (Qiagen, Hilden, Germany) according to the manufacturer’s protocol. The quantity of RNA was assessed via a NanoDrop™ One (Thermo Scientific, Waltham, MA, USA), whereas its integrity was evaluated via the 4200 TapeStation System (Agilent Technologies, Santa Clara, CA, USA). RNA samples with a RIN value of 6.5 or higher were deemed to have acceptable integrity and were subsequently processed for the cDNA library construction.

For the library preparation for RNA sequencing, the KAPA RNA HyperPrep Kit (Roche, Basel, Switzerland) was utilized, following the manufacturer’s instructions. cDNAs were ligated with Unique Dual-Indexed Adapters from the KAPA Kit (Roche, Basel, Switzerland). The concentration of each library was determined via a Qubit fluorometer (Thermo Fisher Scientific, Waltham, MA, USA), and quality was assessed via the 4200 TapeStation System (Agilent Technologies, Santa Clara, CA, USA). The libraries were pooled in equal quantities, and the pooled samples were sent for sequencing at Azenta Genewiz, South Plainfield, NJ, USA. The sequencing was performed on an Illumina NextSeq 500/550 (Illumina, Inc., San Diego, USA) using a 75-cycle run for paired-end reads.

Genes were filtered for at least 75% of the values to be above zero, run through principal component analysis (PCA, with 10 components), and a standard scalar was applied to the PCs.

### 2.7 Proteomics

For the proteomic analysis (protein extraction, sample preparation, mass spectrometry and early data processing), the pond-collected samples were sent to MS Bioworks, Ann Arbor, MI, USA.

For the sample preparation, the cells were washed twice with 1X PBS and lysed in a modified RIPA buffer containing 50 mM Tris HCl (pH 8.0), 150 mM NaCl, 2.0% SDS, 0.1% TX100, and 1X Roche Complete Protease Inhibitor (Roche, Basel, Switzerland). Lysis was performed using 1.4 mm stainless steel beads in a Next Advance Bullet Blender (Next Advance, Troy, NY, USA) for two cycles of 3 minutes each. The samples were then heated to 60 °C for 30 minutes and clarified by centrifugation at 13,200 rpm. Protein was precipitated via trichloroacetic acid (TCA) overnight at −20 °C, after which the pellet was washed and resuspended in a solution of 8 M urea, 50 mM Tris HCl (pH 8.0), and 1X Roche Complete Protease Inhibitor (Roche, Basel, Switzerland). Protein quantification was performed via a Qubit fluorometer (Thermo Fisher Scientific, Waltham, MA, USA), and a pooled sample was created from all the individual samples. For digestion, 2.5 μg of protein was treated overnight with trypsin. The samples were first reduced at room temperature for 1 hour in 12 mM DTT, followed by alkylation for 1 hour at room temperature in 15 mM iodoacetamide. Trypsin was added at an enzyme-to-substrate ratio of 1:20. Each sample was then acidified to 0.3% TFA and subjected to solid-phase extraction (SPE) using a Waters μHLB (Waters, Pleasenton, CA, USA). The SPE process involved activating the matrix with four additions of 500 µL of 70% acetonitrile, equilibrating with four additions of 500 µL of 0.3% TFA, adding the sample, washing the wells with three additions of 500 µL of 0.3% TFA, and eluting in 200 µL followed by an additional 400 µL of 60% acetonitrile in 0.3% TFA. The samples were then frozen at −80 °C and lyophilized overnight.

For the mass spectrometry analysis, 500 ng of each sample was analyzed in triplicate via an LC-MS/MS setup consisting of a Thermo Fisher Vanquish Neo UPLC system (Thermo Fisher Scientific, Waltham, MA, USA) interfaced with a Thermo Fisher Orbitrap Astral (Thermo Fisher Scientific, Waltham, MA, USA). Peptides were loaded onto a trapping column and eluted through a 75 μm analytical column at a flow rate of 350 nL/min, with the column temperature maintained at 55 °C. A 30-minute gradient was used for elution. The mass spectrometer was operated in data-independent acquisition (DIA) mode, capturing full scan MS data at a resolution of 240,000 FWHM over the m/z range of 380-980, followed by 300 sequential 2 m/z precursor isolation windows, with product ions acquired at a resolution of 40,000 FWHM. The maximum ion injection time (IIT) was set to 3.5 ms for DIA, and the normalized collision energy (NCE) was set to 25. An injection of the pooled sample was included at both the beginning and end of the analysis batch.

The initial data processing for the DIA results was performed via DIA-NN (v1.9.1) [29], which facilitated several key functions: conversion of RAW files to the QUANT format, alignment on the basis of retention times, searching data via an in-silico spectral library generated from the FASTA file and iteratively from the spectral library derived from RAW data, filtering of database search results at a 1% false discovery rate (FDR) for both peptides and proteins, calculation of peak areas for detected peptides, and normalization of the data. The data were subjected to percentile normalization for the PCA plots.

### 2.8 Metabolomics

For the metabolomics analysis of the ponds, 1 mL from the samples taken (containing both the media and cells) was sent to Creative Proteomics, Shirley, NY, USA, for untargeted metabolomics analysis. For the blank, samples with only media were taken from the ponds before the algal inoculation. For the sample preparation, the samples were transferred into a tube for lyophilization. Following transfer, 500 µL of 80% methanol was added, and the mixture was vortexed for 30 seconds and then subjected to sonication in an ice-water bath for 30 minutes. The samples were subsequently stored at −20 °C for 1 hour, after which they were vortexed for an additional 30 seconds and centrifuged at 12,000 rpm for 10 minutes at 4 °C, to lyse and remove the cells. From the resulting supernatant, 200 µL was transferred into a vial, and 5 µL of a 0.14 mg/mL solution of DL-o-chlorophenylalanine was added. The final solution was filtered through a 0.22 μm filter in preparation for LC-MS analysis. The separation of the compounds was achieved via the Vanquish Flex UPLC system in conjunction with a Q Exactive Plus mass spectrometer (Thermo Scientific, Waltham, MA, USA), which featured a heated electrospray ionization (ESI) source. The LC separation utilized a Hypersil Gold C18 column (100 × 2.0 mm, 1.9 μm), with the mobile phase consisting of solvent A (0.05% formic acid in water) and solvent B (acetonitrile) under a gradient elution profile: from 0 to 1 minute at 5% B, from 1 to 12.5 minutes increasing from 5% to 95% B, holding at 95% B from 12.5 to 13.5 minutes, returning to 95% B from 13.5 to 13.6 minutes, and finally returning to 5% B from 13.6 to 16 minutes. The flow rate was maintained at 0.3 mL/min, with the column temperature set at 40 °C and the sample manager temperature at 4 °C. The mass spectrometry parameters for both ESI positive (ESI+) and negative (ESI-) modes were configured as follows: for ESI+, the heater temperature was set to 300 °C, the sheath gas flow rate was 45 arb, the auxiliary gas flow rate was 15 arb, the sweep gas flow rate was 1 arb, the spray voltage was 3.0 kV, the capillary temperature was 350 °C, and the S-Lens RF level was 30%. For ESI-, the heater temperature remained at 300 °C, with the same sheath and auxiliary gas flow rates as those used for ESI+, but the spray voltage was adjusted to 3.2 kV, and the S-Lens RF level was increased to 60%. For the bioinformatic analysis (see section 2.9), we considered the metabolites whose abundance was more than 10x greater than the abundance of the metabolites in the blank to be significant.

For the small-scale metabolomics, 100 mL cultures of *M.minutum* were grown in 250 mL Erlenmeyer flasks (in triplicate) under the same conditions and media described in 2.1. On day 5 the cultures have been 2/3 diluted and FDO1 [19] [18] was added at a 0.05% volume concentration. Samples of 5ml from each flask were collected on days 0, 3, 4, 5, 7, 10, 11, 12, 17 and 18. The LC-MS and bioinformatic analysis were conducted as described for the pond samples.

### 2.9 Bioinformatics

Principal component analysis (PCA) was performed in R with the factoextra package [30]. For 3D PCA analysis, the first 3 components were plotted for each replicate per day, and the centroid point was also plotted on the same axes. Day-specific standard deviation from the centroid (based on replicates) is shown by a shaded sphere.

For RNA sequencing analysis, the following sequence processing steps and associated packages were used: the fastq file was quality-checked with FastQC, and adapter trimming was performed with Trimmomatic [31], t The pseudoalignment of the transcripts to the transcriptome of *M. minutum* (as it was published and publicly available by [17]) and the quantification of their abundance were performed with the Kallisto package [32]. The integration of the Kallisto data and the differential gene expression analysis, including the time course analysis, was performed with Sleuth [33]. The GEMMA package [34] was used for the correlation of the gene expression with specified quantitative phenotypes.

Additional tools for the multi-omic analyses include ggplot2 [35] for graphic representation and pheatmap [36] for the heatmap.

De novo motif enrichment analysis of the promoter regions (200 bps upstream of ATG) was performed with the Homer Motif Discovery and Analysis software [37].

We used the Data Integration Analysis for Biomarker Discovery using Latent variable approaches for Omics studies (DIABLO) framework from the R software package mixOmics (version 6.28.0) to combine the transcriptomics, proteomics, and metabolomics datasets [38]. Specifically, we compared these data across Day 0 and Day 1 and subsequently Day 4 and Day 5 to identify key features of culture scale up and infection, respectively. The metabolomics and proteomics datasets were log transformed prior to this analysis, and the transcriptomics data were normalized to transcripts per million (TPM) values. The proteomic and transcriptomic datasets were further filtered via the nearZeroVar function from the caret R package (version 7.0-1), with a unique cutoff of 20 for the proteomics dataset and selecting transcripts with variances of greater than or equal to 200 were selected. In the DIABLO design matrix, we prioritized correlations between the three omics datasets by choosing a 0.6 weighted model. The features from latent component 1, from the Day 0 and Day 1 comparison and from the Day 4 and Day 5 comparison were then compiled for comparison across all four days via the R package pheatmap (version). The values were scaled and centered prior to plotting, and both rows and columns were clustered via Euclidean distance.

For the correlation analysis, genes and proteins were filtered for members that are expressed in at least 50% of samples, and at least 2/3 of replicates. Then samples were percentile normalized, and averaged across replicates. Following this, a pairwise Pearson correlation was conducted, and the results were visualized in a heat map.

For the protein-protein interaction networks, STRING (https://string-db.org/) was used to construct protein-protein interaction (PPI) networks. For the scale-up stress (D0-D1) and the amoeboaphelid stress (D4-D5), changing genes and proteins were combined into one dataset. As there are no annotations on STRING for *Monoraphidium minutum*, we used the *Chlamydomonas reinhardtii* annotation to find the closest match, and submitted those matches to STRING to generate PPI networks. The colorimetric indication of the nature of each protein-protein interaction is being explained in details at https://string-db.org/.

Based on the scant ontology analysis of *Monoraphidium minutum* and based on a recent paper [39] stating that GPT4 is well-suited to ontology analysis, we prompted GPT4 as follows to identify cohesive pathways from gene descriptions: “Given the gene functions from this prompt and the previous one, give me a list of plausible, cohesive pathways that are being regulated in a green algae *(Monoraphidium minutum*). Highlight the similarities and differences in the pathways that are engaged in the two lists.” We also independently verified claims with references, knowing that GPT4 and other LLMs are known to hallucinate references.

## 3. Results

### 3.1 Phenotypic and conditional measurements reveal distinct acclimation, growth and infection dynamics in algae in open pond systems

We performed a longitudinal analysis of algal growth in two 1000 L open raceway ponds, and introduced an amoeboaphelid pathogen on day 4 of the experiment, tracking a total of 12 variables. Phenotypic measurements included cell count, AFDW, chlorophyll concentration, total carbohydrate content, total H3 histone concentration, cell size, and caspase activity (Additional File 1). In addition, we measured five environmental variables within the ponds, including %DO, pH, conductivity, temperature, and light intensity.

Across the time-course, we observed 3 growth phases of interest (color-coded in Fig. 2A):

a. The acclimation or lag phase (cyan). This phase occurred immediately after the inoculation in the open system with refreshed nutrients. This period lasted 2 days, and growth during this phase was slower than that during the other phases.
b. b). The log phase (yellow). The cells were fully acclimated to the open-pond conditions and grown exponentially. This phase lasted for 4 days before the dilution and the infection and 3 days after dilution.
c. Crash phase (brown). This phase is caused by algal death, which leads to the complete overtake of the ponds by the pathogen and other opportunistic species.

**Fig. 2:**
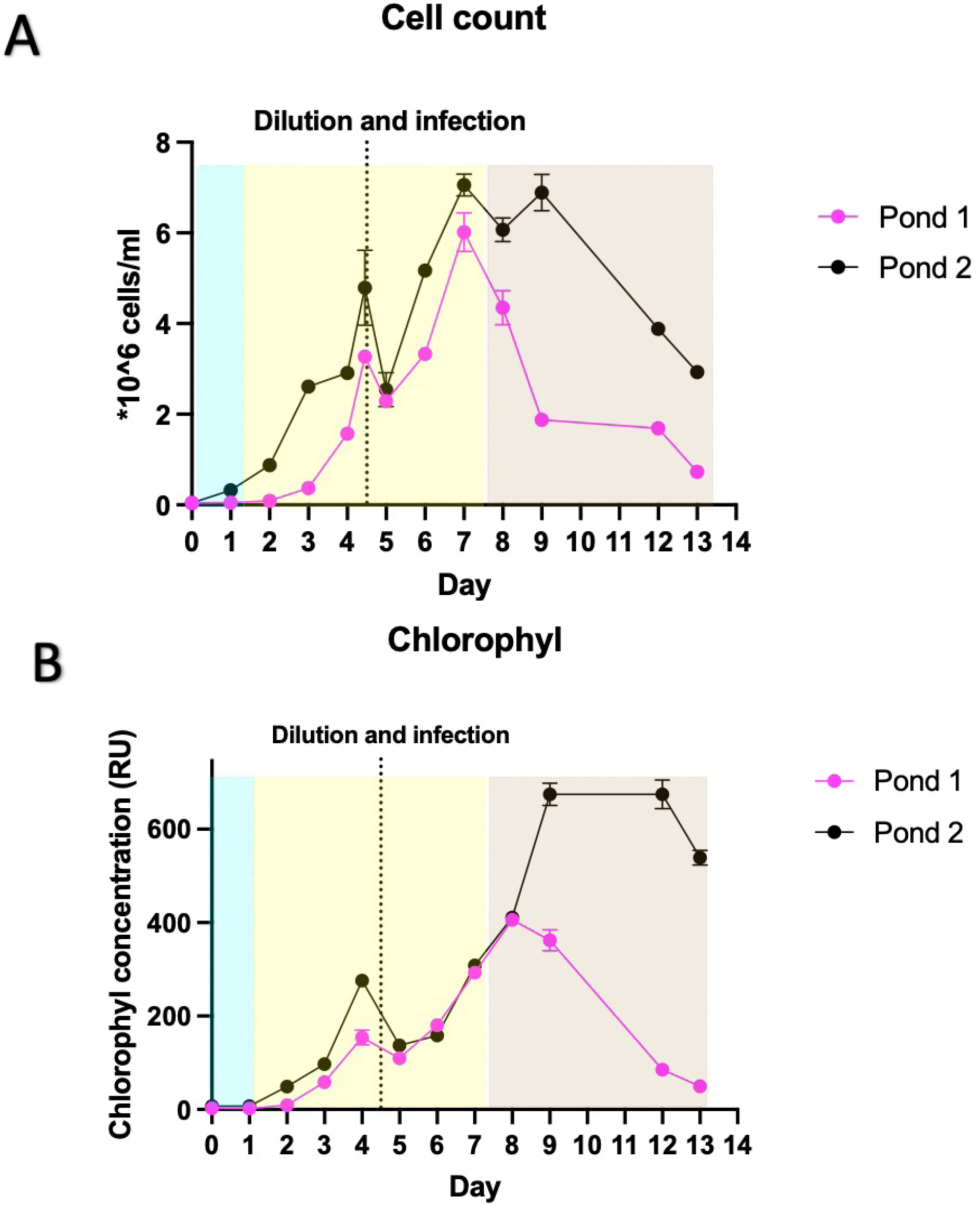
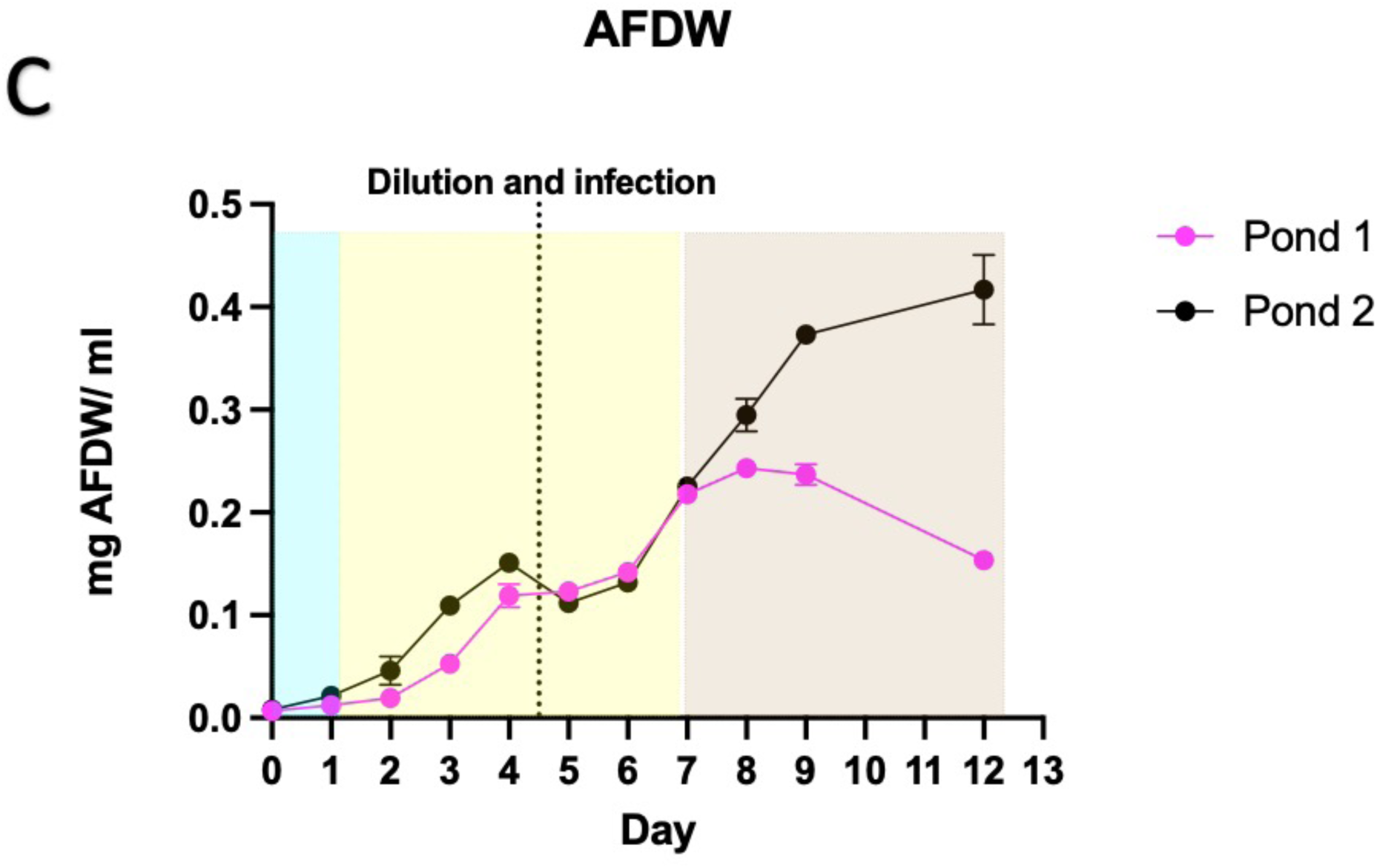
Growth tracking of the M. minutum pond cultures. A) The cell count between the three ponds. B) Chlorophyl concentration in relative units (RU) from florescence measurements. C) The Ash-free dry weight (AFDW) of algae cells in mg per mL of culture.

The ponds behave largely similarly, although we do observe a divergence of phenotypes at later stages of infection, as documented in Fig. 2.

To understand whether stress conditions trigger programmed cell death, we measured caspase activity throughout the experiment. Caspases are proteases that catalyze the breakdown of proteins during programmed cell death [40]. We observed that caspase activity initially increased after pond inoculation and then gradually decreased as the cells acclimated to their new environment (Additional File. 2). Notably, we did not observe an increase in caspase activity after infection with amoeboaphelid, suggesting that the reduction in the algal cell count could be attributed to alternative mechanisms, perhaps uncoupled from programmed cell death. To identify a specific pathway responsible, more molecular studies focused on underlying mechanism by which cells become depleted are needed. [41, 42]

For an integrated view of the phenotypic and conditional profile of the longitudinal experiment, principal component analysis (PCA) was performed with the 12 phenotypic and environmental variables. Dimension 1 represented 49.4% of the explained variance (Fig. 3A) and separated the variables that are indicative of growth, from those indicative of stress. This provides insight into the competing pressures that the algae face during cultivation, and the contributions of these pressures to outcomes.

**Fig. 3:**
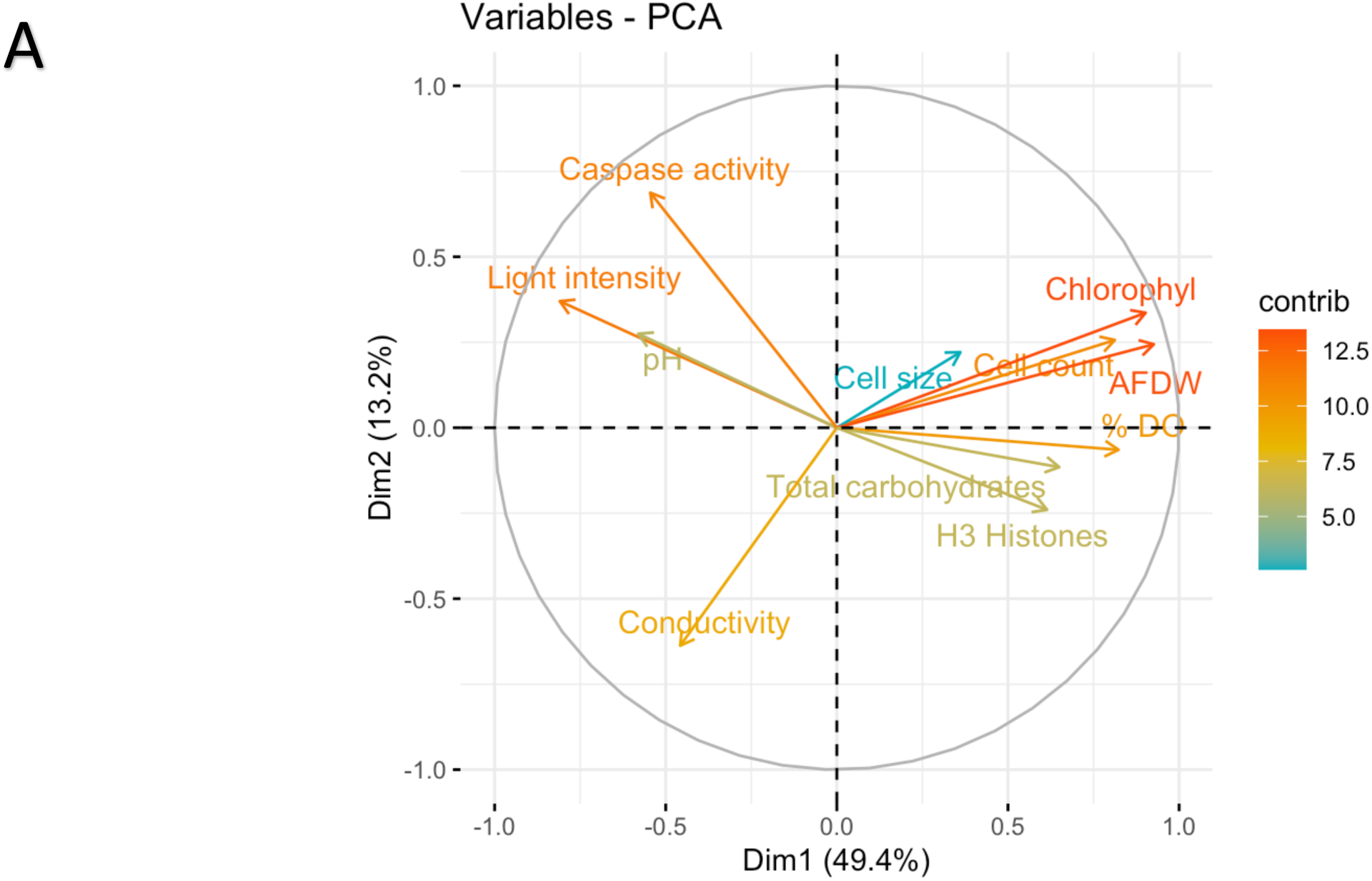
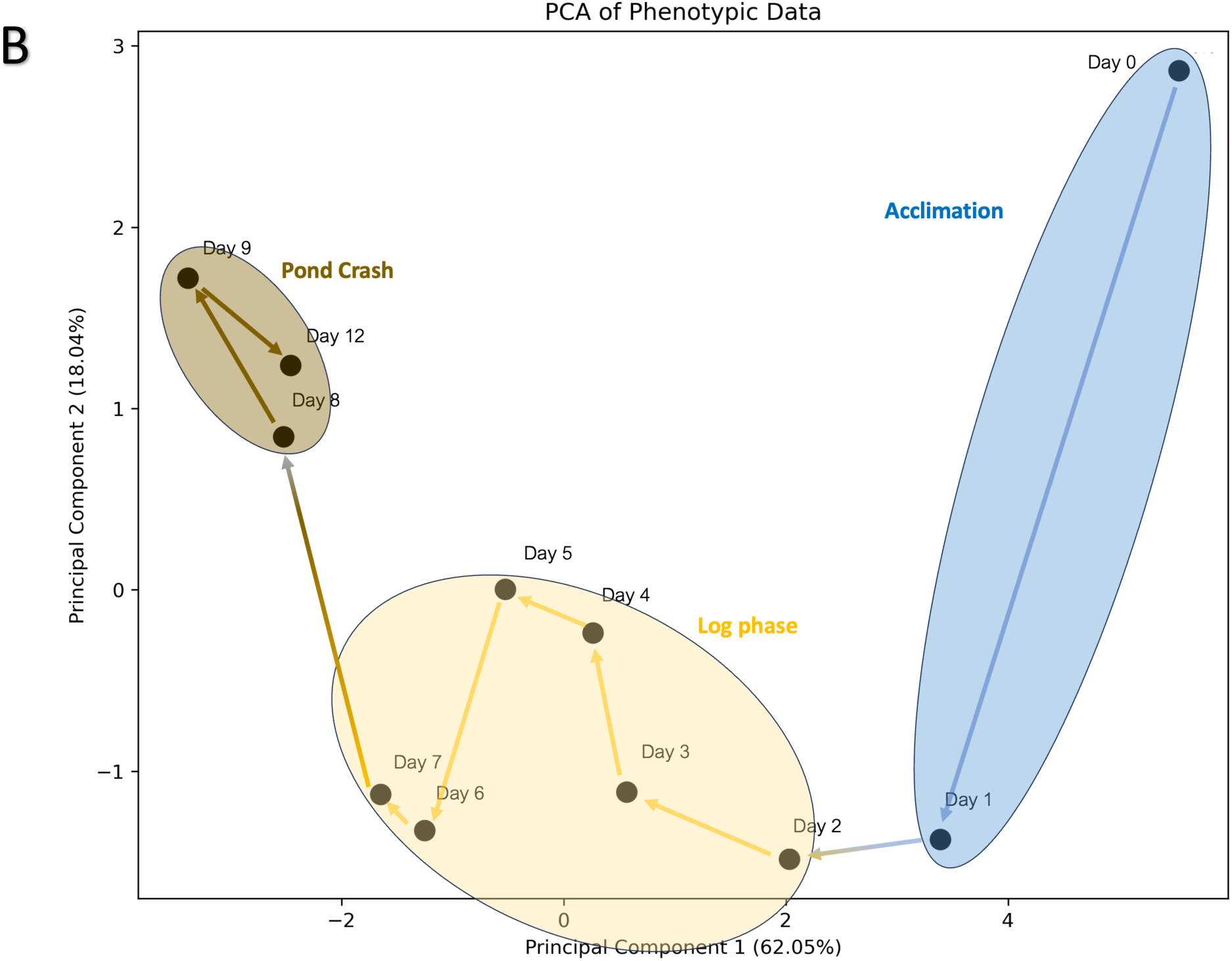
PCA for 12 variables as measured at the two ponds. A) PCA for the 12 variables used. The color of each line represents the contribution of each variable to the principal axes. Created with factoextra in R. B) Each day of the cultivation, plotted as individuals. The arrows are indicating the trajectory of each day during the culture run.

We observe that the three distinct phases (acclimation, log, and crash) are resolved away from each other in the PCA biplot shown in Figure 3B, and we can discern a trajectory in PC space as the longitudinal experiment progresses, which is shown by connecting temporally adjacent points. While the largest adjacent transition occurred during scale-up (Day 0 to Day 1), we also observed a significant shift upon infection into a distinct PC space highlighted in brown in Figure 3B. This visualization further emphasizes the distinction between the conditions and outcomes during these phases and suggests the presence of distinct molecular profiles within the algae that likely precede outcomes. To this end, we performed transcriptomic, proteomic and metabolomic analyses across the first 6 days of this experiment.

Community composition analysis though 16/18S was also performed in order to determine the eukaryotic and bacterial composition of the ponds throughout the culture run (Fig. 4, Additional File 2). Those systems are open and expected to contain a wide variety of organisms growing in them other that *M.minutum* and the pathogen we introduced. The bacterial composition (Fig.4A), possibly consisting of both environmental and algae-associated bacteria, seems relatively stable throughout the run, with the exception of day 19, which goes beyond the total crash of the cultures. From the eukaryotic composition (Fig.4B) we can verify that, while *M. minutum* is predominant in the ponds throughout the duration of the experiment, as expected, other green algal species were co-present in relatively far smaller proportions. Notably, the pathogen was first detected on day 12, despite the fact that the culture was impacted by its presence much earlier, showing that alternative biomarkers are required for early detection of infection.

**Fig. 4:**
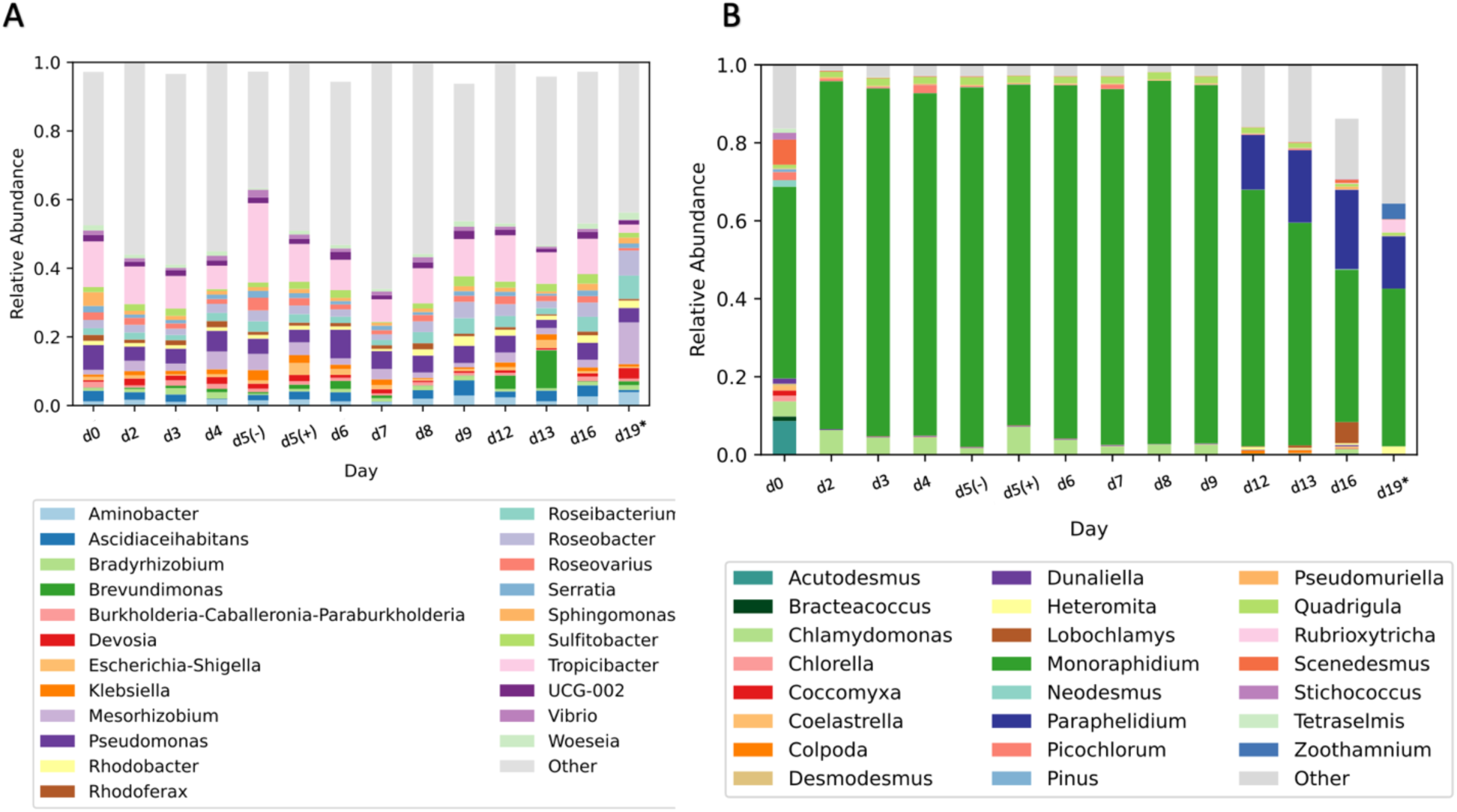
Microbial community composition of the ponds throughout the run until after the complete pond crashes. A) Bacterial genera. B) Eukaryotic genera.

### 3.2 Transcriptomic stability tracks with optimal growth and culture resiliency

With the goal of identifying molecular patterns within the algae that precede the observed phenotypic outcomes, we performed transcriptomic analysis from Day 0 to Day 6 (Additional File 3). The samples thus represented the acclimation period (day 0 and day 1), the preinfection log phase (day 2- day 4) and the early infection days (days 5 and 6). Throughout this period the culture is in continuous log phase and while we acknowledge that the growth and density can have an impact on the molecular state of the algae, we perform the infection prior to the slow-down and plateau phase to mitigate these effects.). For the scale-up, part of the stressor is the new density the algae are faced with, so we do not specifically adjust for this. A considerable percentage of transcripts (12.3%) were significantly differentially expressed from day 0 (the inoculation day) to day 1, reflecting the transcriptional acclimation to the scale transition from the 20 L carboys to the open 1000 L ponds (Fig.5A). This difference in expression was multiple times greater than any daily difference observed during the following days. This included days after infection where only 0.7% of the transcripts were significantly differentially expressed. Consistent with the algae being in a homeostatic growth phase, we saw very few differences in gene expression across non-stressor days, suggesting that the majority of changes occur as a result of stress rather than growth.

**Fig. 5:**
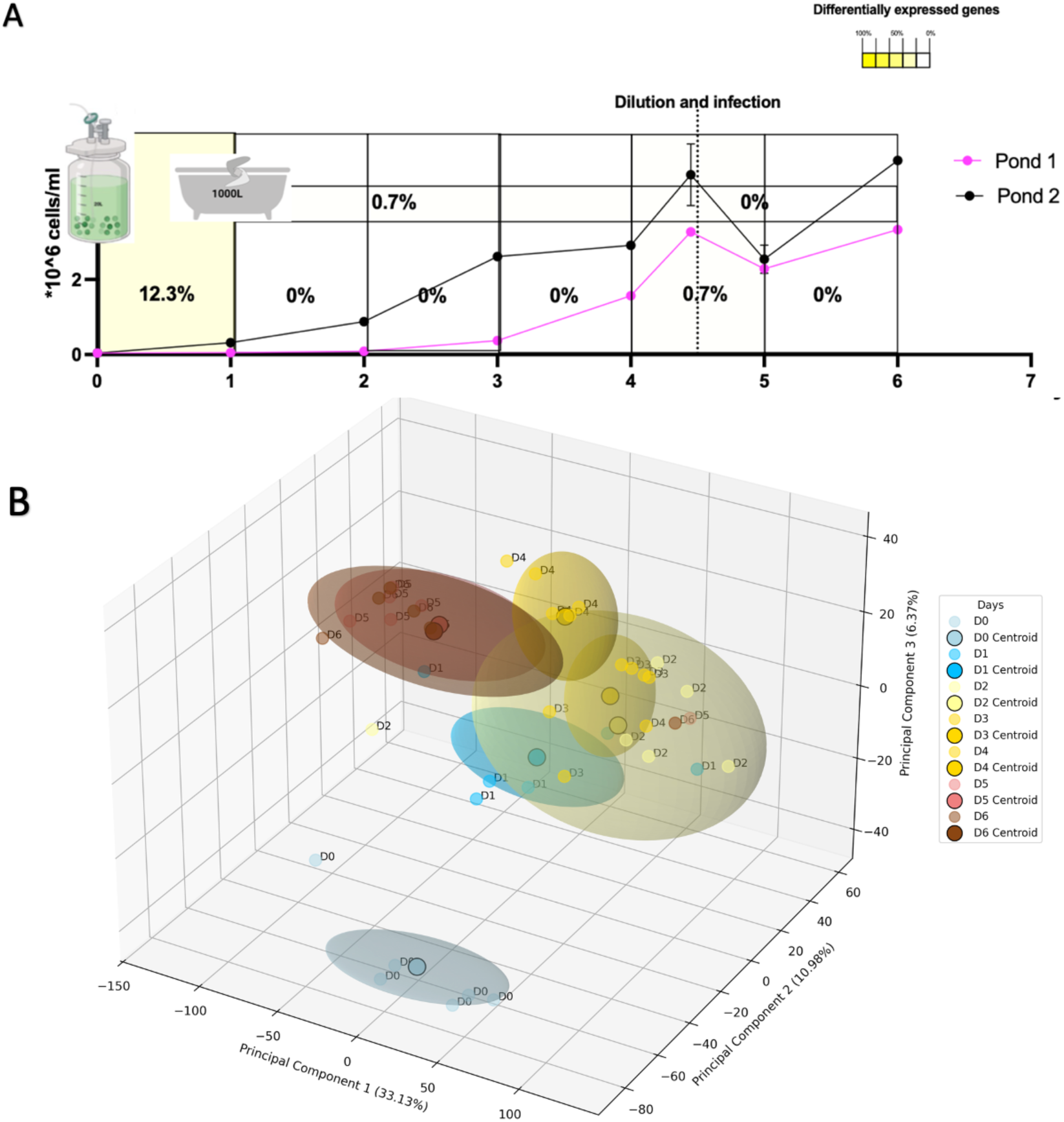
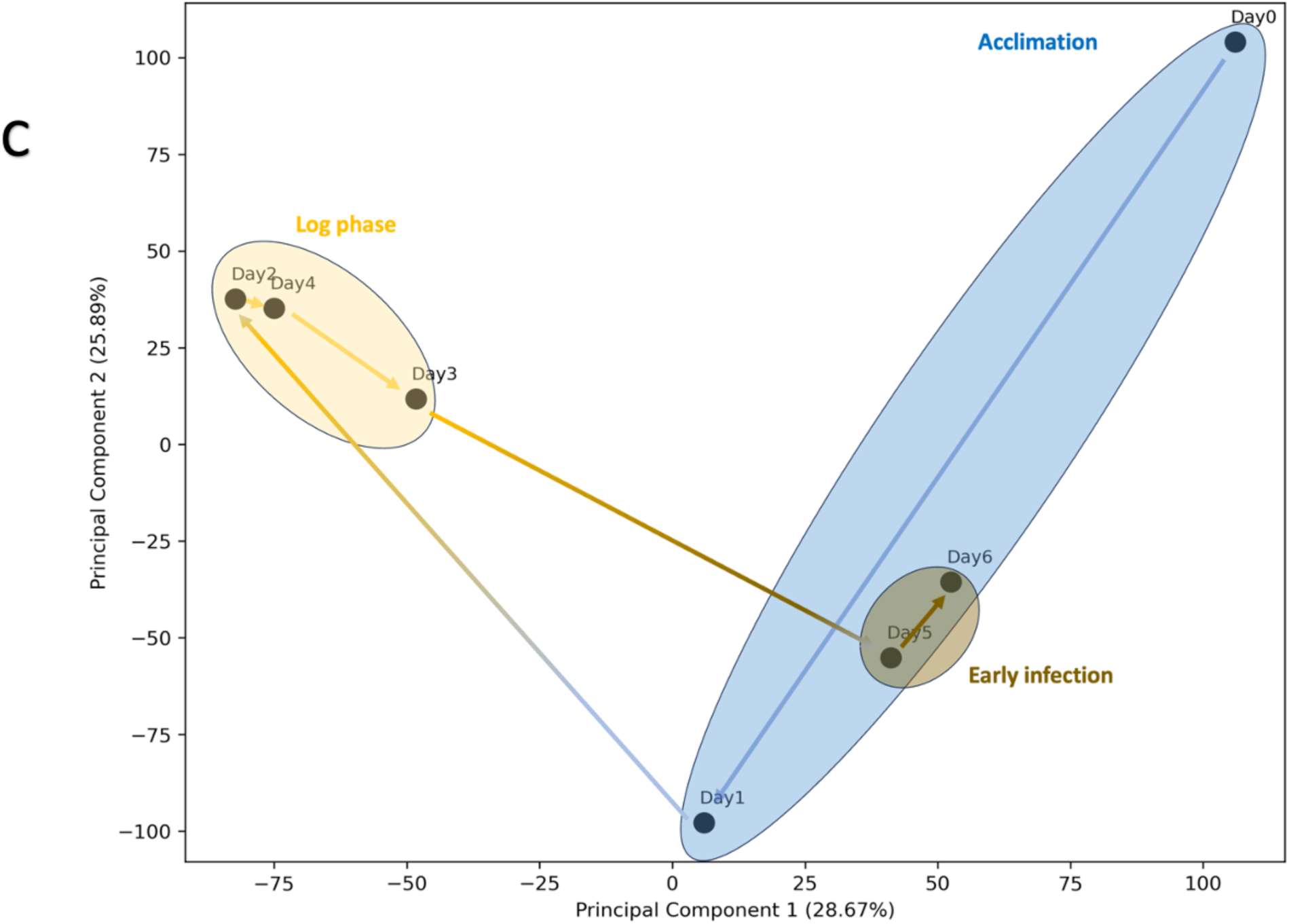
Comparative analysis of the transcript abundances overlaid with cell growth results. A) Growth curve of both ponds over the time course, with transcriptomic data collected on the same day as growth was measured (as shown by points on the curve). Percentages of differentially expressed genes between specific daily points within the cultivation are shown by the numbers. B) 3D PCA analysis of the transcriptomics, showing the distribution of all the technical replicates for each day from the two ponds. Centroids are indicated with a darker point, and standard deviation is indicated by the shaded circle. C) 2D representation of the PCA showing the average points for each day. The arrows indicate the trajectory of the time course.

We performed PCA on the transcriptomic data to determine whether the longitudinal trajectory was comparable to that of our phenotypic and environmental variables (Fig. 5B and 5C). The transition from Day 0 to Day 1 was the most striking, reflecting the rapid and large transcriptomic response to the 1000 L pond conditions. The distributions of the days corresponding to the pre-infection log phase, on the other hand, are closer in PC space, reflecting transcriptional stability during that growing stage. A clear distinction between the distribution of the preinfection and postinfection days was also observed, as a result of both the culture dilution and the introduction of the pathogen. In fact, unlike in Fig. 3B where the measured environmental and phenotypic variables postinfection were in a distinct cluster away from the scale-up, the early postinfection transcriptomic profile trends back toward the acclimation phase profile but still remains distinct from them.

The annotation of *M. minutum* genome is incomplete, making traditional pathway analysis complicated. To further investigate the regulation behind those transcriptional changes we performed a motif analysis among the promoter regions of the genes that were significantly differentially expressed to identify regions that could be bound by conserved transcription factors. We focused on the transitions between day 0 and day 1 (Table 1), and between day 4 and day 5 (Table 2). The ten most enriched motifs that emerged from each group are shown in Table 1 and Table 2, alongside their best matches from other organisms. Most identified motifs are unique to each transition.

**Table 1:**
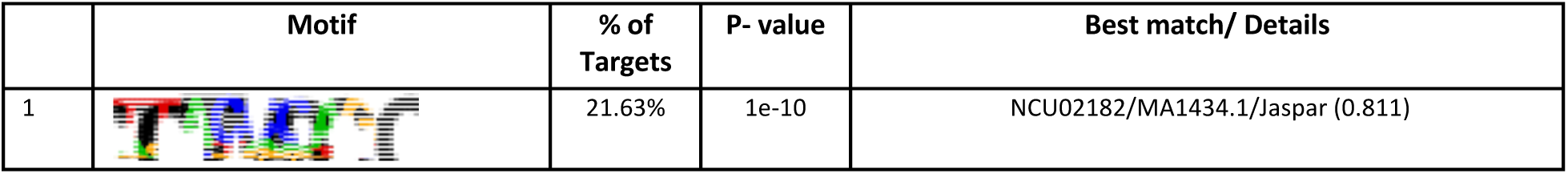
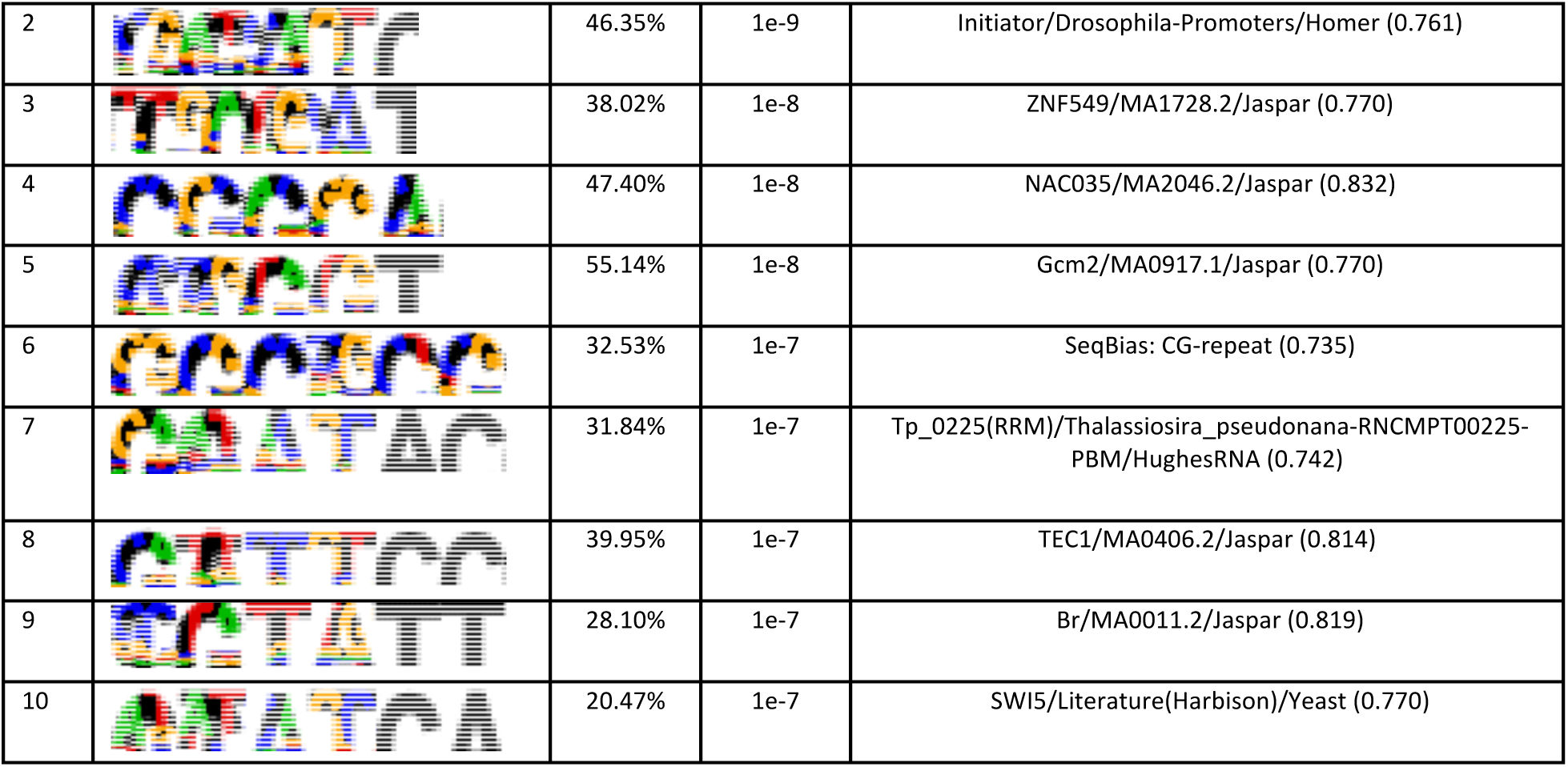
Enriched promoter motifs at the scale adaptation-related genes.

|  | Motif | % of Targets | P- value | Best match/ Details |
| --- | --- | --- | --- | --- |
| 1 |  | 21.63% | 1e-10 | NCU02182/MA1434.1/Jaspar (0.811) |
| 2 |  | 46.35% | 1e-9 | Initiator/Drosophila-Promoters/Homer (0.761) |
| 3 |  | 38.02% | 1e-8 | ZNF549/MA1728.2/Jaspar (0.770) |
| 4 |  | 47.40% | 1e-8 | NAC035/MA2046.2/Jaspar (0.832) |
| 5 |  | 55.14% | 1e-8 | Gcm2/MA0917.1/Jaspar (0.770) |
| 6 |  | 32.53% | 1e-7 | SeqBias: CG-repeat (0.735) |
| 7 |  | 31.84% | 1e-7 | Tp_0225(RRM)/Thalassiosira_pseudonana-RNCMPT00225-PBM/HughesRNA (0.742) |
| 8 |  | 39.95% | 1e-7 | TEC1/MA0406.2/Jaspar (0.814) |
| 9 |  | 28.10% | 1e-7 | Br/MA0011.2/Jaspar (0.819) |
| 10 |  | 20.47% | 1e-7 | SWI5/Literature(Harbison)/Yeast (0.770) |

**Table 2:**
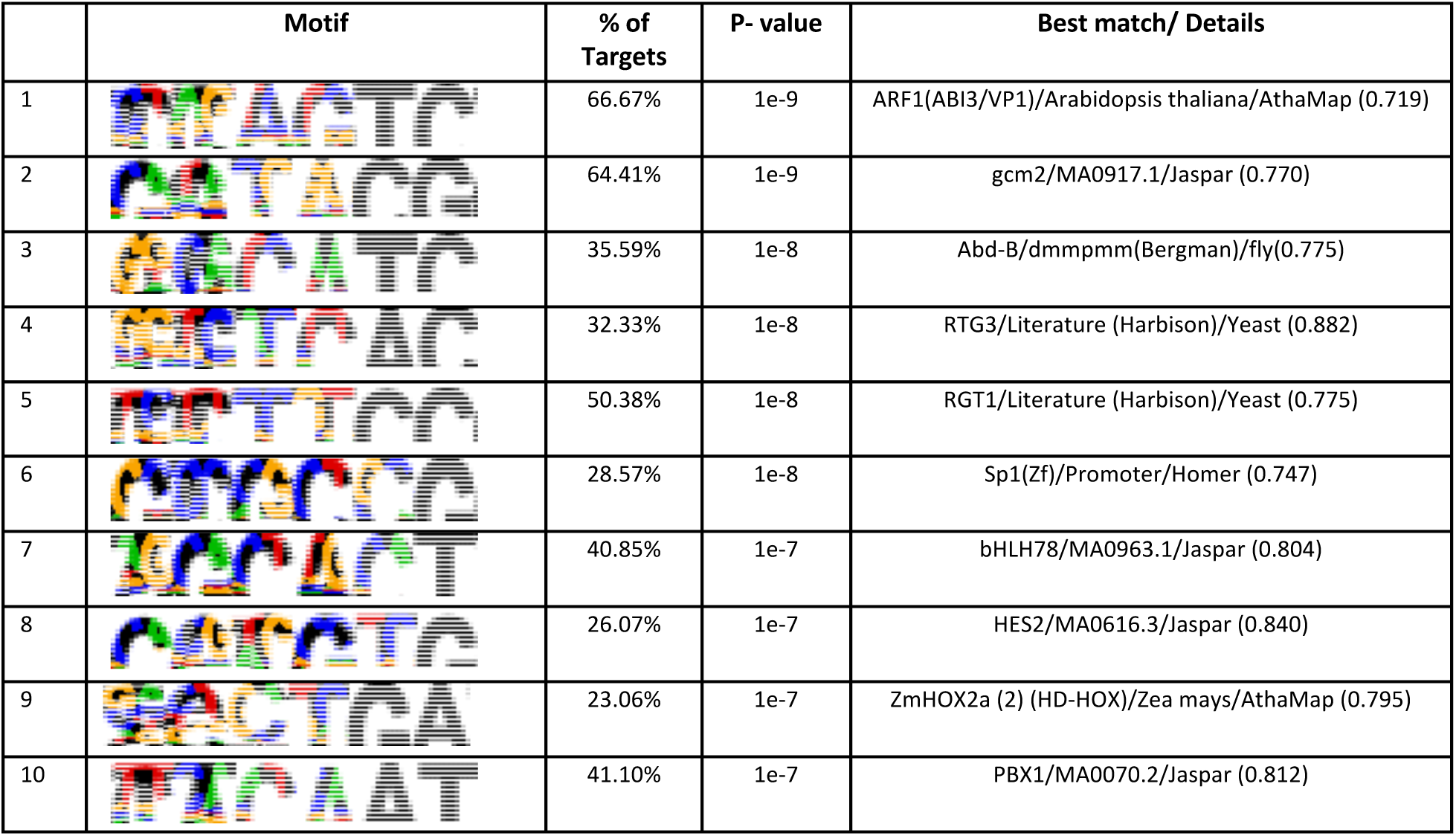
Enriched promoter motifs at the infection-related genes.

|  | Motif | % of Targets | P- value | Best match/ Details |
| --- | --- | --- | --- | --- |
| 1 |  | 66.67% | 1e-9 | ARF1(ABI3/VP1)/Arabidopsis thaliana/AthaMap (0.719) |
| 2 |  | 64.41% | 1e-9 | gcm2/MA0917.1/Jaspar (0.770) |
| 3 |  | 35.59% | 1e-8 | Abd-B/dmmpmm(Bergman)/fly(0.775) |
| 4 |  | 32.33% | 1e-8 | RTG3/Literature (Harbison)/Yeast (0.882) |
| 5 |  | 50.38% | 1e-8 | RGT1/Literature (Harbison)/Yeast (0.775) |
| 6 |  | 28.57% | 1e-8 | Sp1(Zf)/Promoter/Homer (0.747) |
| 7 |  | 40.85% | 1e-7 | bHLH78/MA0963.1/Jaspar (0.804) |
| 8 |  | 26.07% | 1e-7 | HES2/MA0616.3/Jaspar (0.840) |
| 9 |  | 23.06% | 1e-7 | ZmHOX2a (2) (HD-HOX)/Zea mays/AthaMap (0.795) |
| 10 |  | 41.10% | 1e-7 | PBX1/MA0070.2/Jaspar (0.812) |

**Table 3:** The annotations of the top 5 genes whose expression was upregulated with light.

| JGI ID | beta | Gene product |
| --- | --- | --- |
| 427491 | 486.8095 | ribosomal protein S9 component of cytosolic 80S ribosome and 40S small subunit |
| 412255 | 479.0458 | 40S ribosomal protein S29-like |
| 420537 | 442.9904 | putative 40S ribosomal protein S3a |
| 431691 | 419.257 | Translation protein SH3-like domain-containing protein |
| 420483 | 377.4611 | component of cytosolic 80S ribosome and 60S large subunit |

**Table 4:** The annotations of the top 5 genes whose expression was downregulated with light.

| JGI ID | beta | Gene product |
| --- | --- | --- |
| 518027 | -257.7188 | heat shock cognate 70 kDa protein 4-like |
| 474353 | -195.5336 | hypothetical protein |
| 453947 | -167.1721 | eukaryotic translation initiation factor 5 |
| 437942 | -161.7026 | photosystem I protein P30 |
| 272372 | -155.3597 | hypothetical protein |

### 3.3 Transcriptomic and phenotypic data reveal the light and dissolved oxygen (DO) regulation of protein translation and photosynthesis

One of the advantages of performing time-course transcriptomics on large open cultures in a poorly studied species such as *M. minutum* is that it allows for the extraction of reliable and useful information about specific environmentally triggered expressed genes. Light intensity and DO are two environmental factors known to influence outcomes in algae. To understand their role in the outcome of our cultures, we used a program called GEMMA that applies mixed linear models to connect phenotypes with large-scale molecular data [34]. We identified transcripts whose expression was positively or negatively correlated with light intensity and %DO across samples (Figs. 6 and 7).

**Fig. 6:**
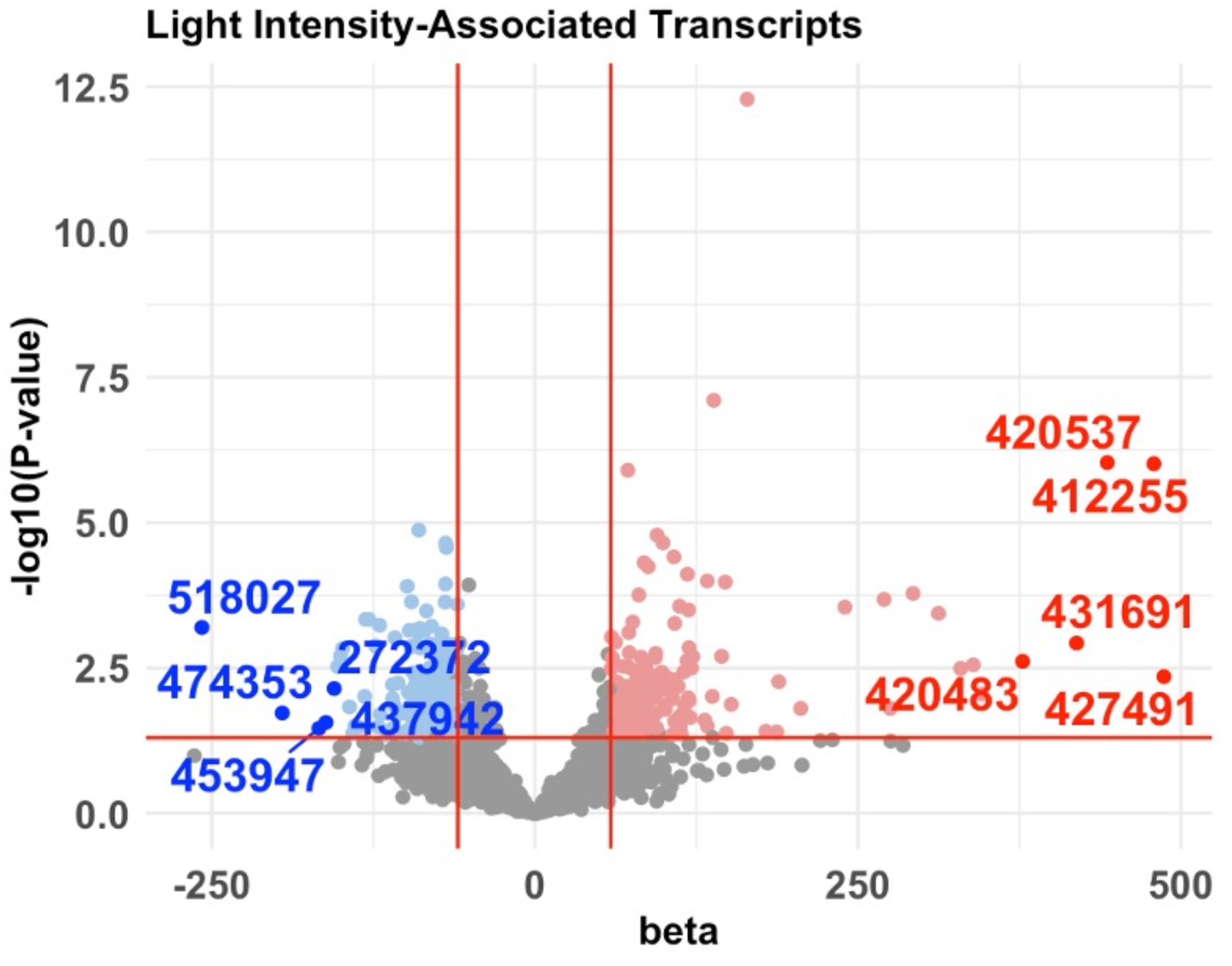
GEMMA analysis, revealing the genes which are correlated with light intensity. The volcano plot shows the significance and the extent of the correlation between the light intensity and the transcript abundance. The top 5 up- and downregulated genes by light are indicated.

**Fig. 7:**
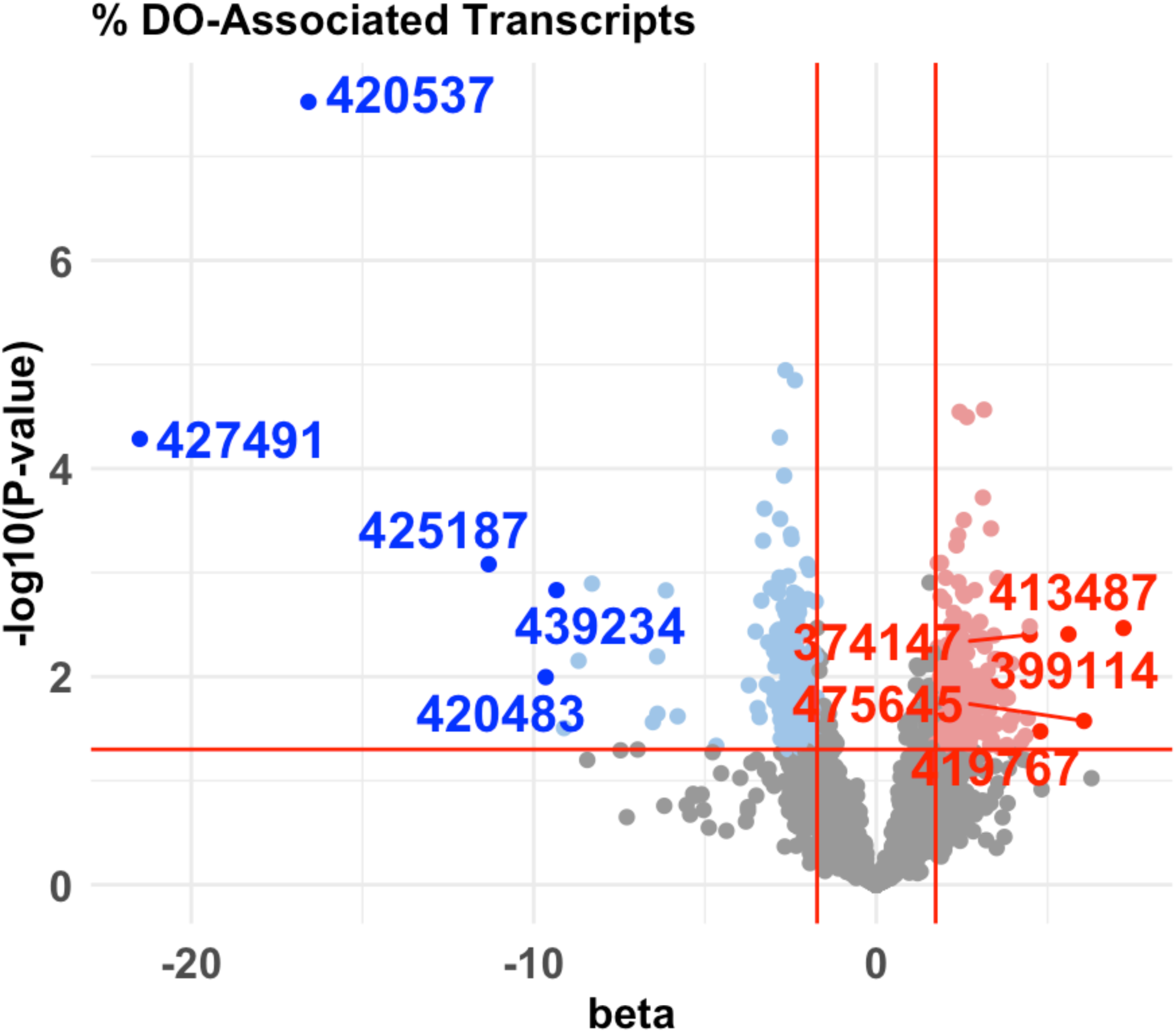
GEMMA analysis, revealing genes correlated with %DO. The volcano plot shows the significance and the extent of the correlation between the light intensity and the transcript abundance. The top 5 up- and downregulated genes by %DO are indicated.

[43]To further understand the regulatory genomic elements that might contribute to light-dependent changes in gene expression, we performed a motif analysis on the promoter regions of all the genes whose expression was significantly up- or downregulated by light (Tables 5 and 6). Among the most significant motifs from the upregulated genes, two corresponded to transcription factors from two higher plant species, *Arabidopsis* and *Zea mays*.

**Table 5:**
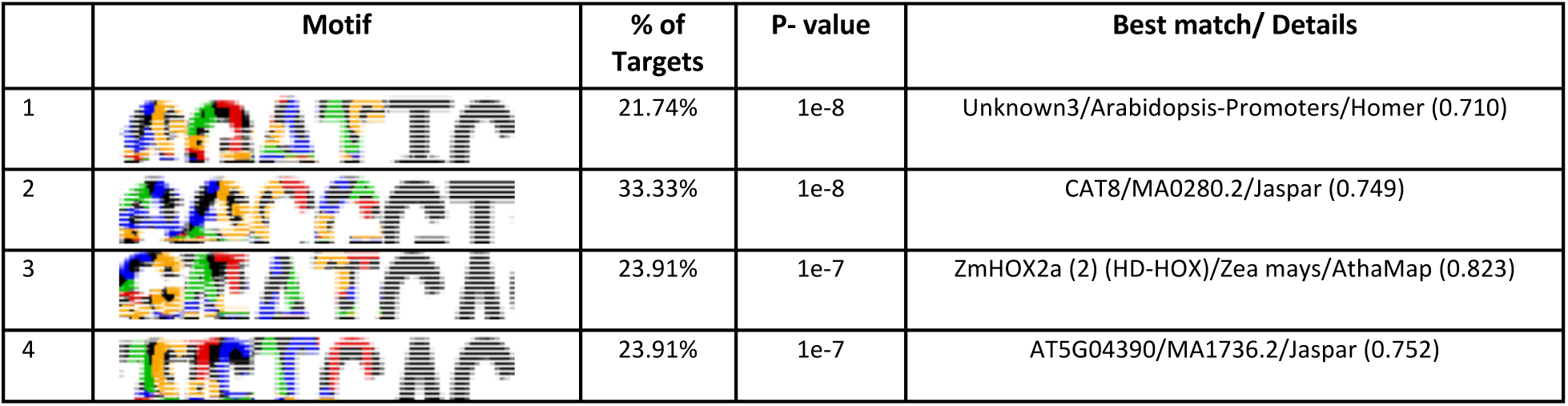
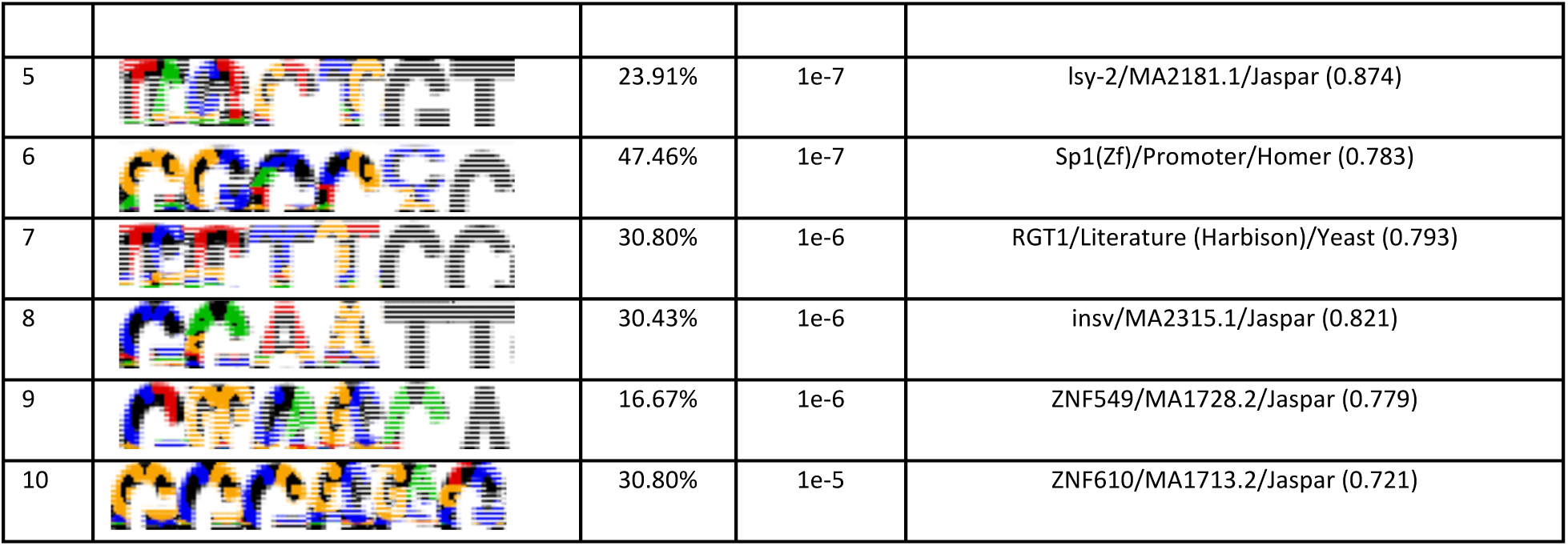
Enriched promoter motifs at the light upregulated genes.

|  | Motif | % of Targets | P- value | Best match/ Details |
| --- | --- | --- | --- | --- |
| 1 |  | 21.74% | 1e-8 | Unknown3/Arabidopsis-Promoters/Homer (0.710) |
| 2 |  | 33.33% | 1e-8 | CAT8/MA0280.2/Jaspar (0.749) |
| 3 |  | 23.91% | 1e-7 | ZmHOX2a (2) (HD-HOX)/Zea mays/AthaMap (0.823) |
| 4 |  | 23.91% | 1e-7 | AT5G04390/MA1736.2/Jaspar (0.752) |
| 5 |  | 23.91% | 1e-7 | Isy-2/MA2181.1/Jaspar (0.874) |
| 6 |  | 47.46% | 1e-7 | Sp1(Zf)/Promoter/Homer (0.783) |
| 7 |  | 30.80% | 1e-6 | RGT1/Literature (Harbison)/Yeast (0.793) |
| 8 |  | 30.43% | 1e-6 | insv/MA2315.1/Jaspar (0.821) |
| 9 |  | 16.67% | 1e-6 | ZNF549/MA1728.2/Jaspar (0.779) |
| 10 |  | 30.80% | 1e-5 | ZNF610/MA1713.2/Jaspar (0.721) |

**Table 6:**
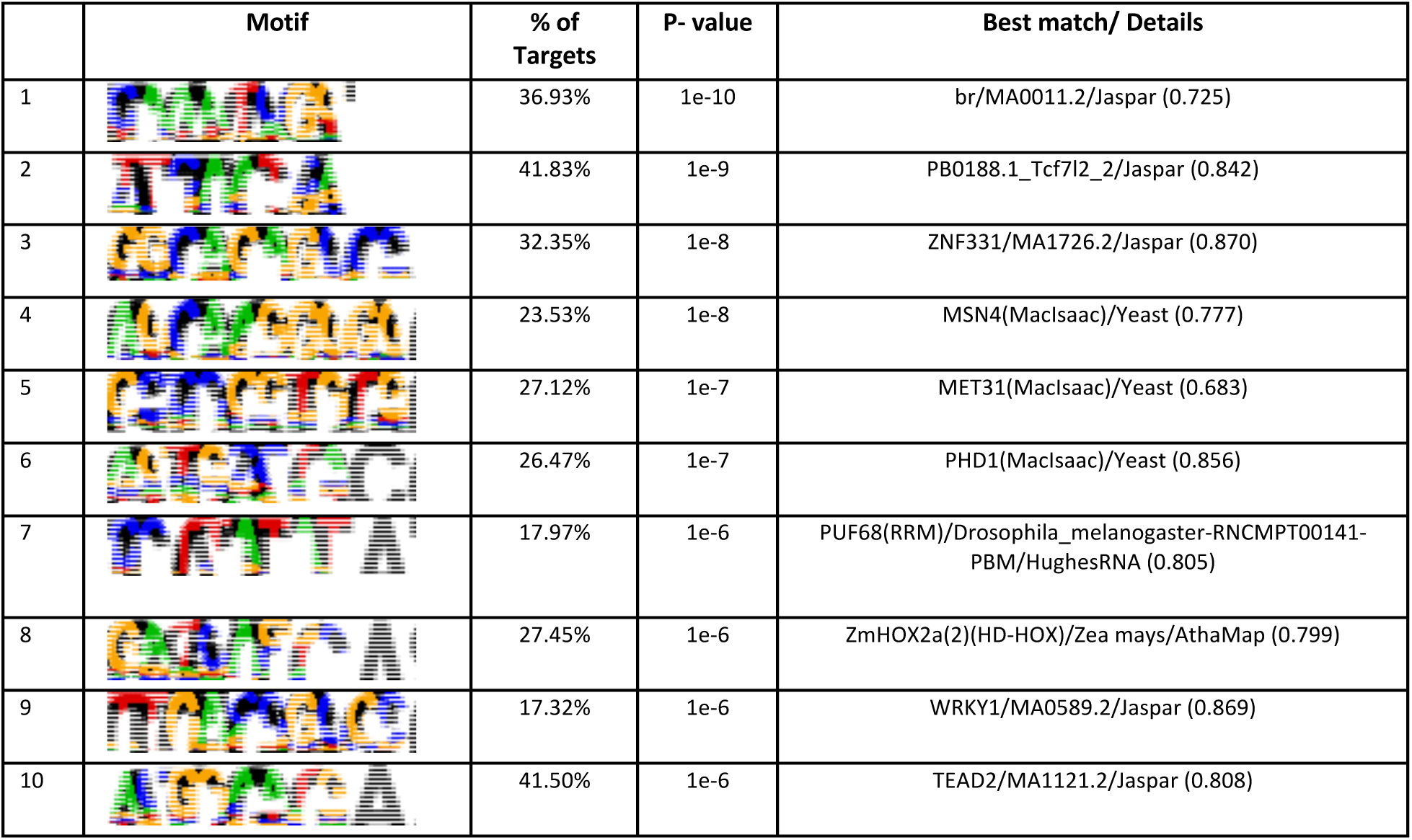
Enriched promoter motifs at the light downregulated genes.

|  | Motif | % of Targets | P- value | Best match/ Details |
| --- | --- | --- | --- | --- |
| 1 |  | 36.93% | 1e-10 | br/MA0011.2/Jaspar (0.725) |
| 2 |  | 41.83% | 1e-9 | PB0188.1_Tcf7l2_2/Jaspar (0.842) |
| 3 |  | 32.35% | 1e-8 | ZNF331/MA1726.2/Jaspar (0.870) |
| 4 |  | 23.53% | 1e-8 | MSN4(MacIsaac)/Yeast (0.777) |
| 5 |  | 27.12% | 1e-7 | MET31(MacIsaac)/Yeast (0.683) |
| 6 |  | 26.47% | 1e-7 | PHD1(MacIsaac)/Yeast (0.856) |
| 7 |  | 17.97% | 1e-6 | PUF68(RRM)/Drosophila_melanogaster-RNCMPT00141-PBM/HughesRNA (0.805) |
| 8 |  | 27.45% | 1e-6 | ZmHOX2a(2)(HD-HOX)/Zea mays/AthaMap (0.799) |
| 9 |  | 17.32% | 1e-6 | WRKY1/MA0589.2/Jaspar (0.869) |
| 10 |  | 41.50% | 1e-6 | TEAD2/MA1121.2/Jaspar (0.808) |

A similar GEMMA analysis was repeated with the genes most correlated with %DO. Among the ten most upregulated and downregulated genes correlated with %DO (Fig. 7A, Tables 7 and 8), multiple genes related to photosynthesis and carbon fixation were identified. A phosphoribulokinase/uridine kinase, an enzyme responsible for a key step in the Calvin-Benson cycle, a light-harvesting protein of Photosystem I, and a protein involved in programmed cell death were among the most upregulated genes correlated with DO. We performed motif analysis on the promoter regions of all the genes correlated with DO (Tables 9 and 10). We observed enrichment of the binding site for the same transcription factor from the diatom *Thalassiosira pseudonana* at both the up- and downregulated genes from the DO.

**Table 7:** The annotations of the top 5 genes whose expression was upregulated with %DO.

| JGI ID | beta | Gene product |
| --- | --- | --- |
| 413487 | 7.216919 | Phosphoribulokinase / Uridine kinase family-domain containing protein |
| 475645 | 6.064846 | hypothetical protein |
| 399114 | 5.610648 | expressed protein |
| 419767 | 4.78923 | light-harvesting chlorophyll-a/b protein of photosystem I |
| 374147 | 4.481988 | Programmed cell death protein 2, C-terminal putative domain-domain containing protein |

**Table 8:**
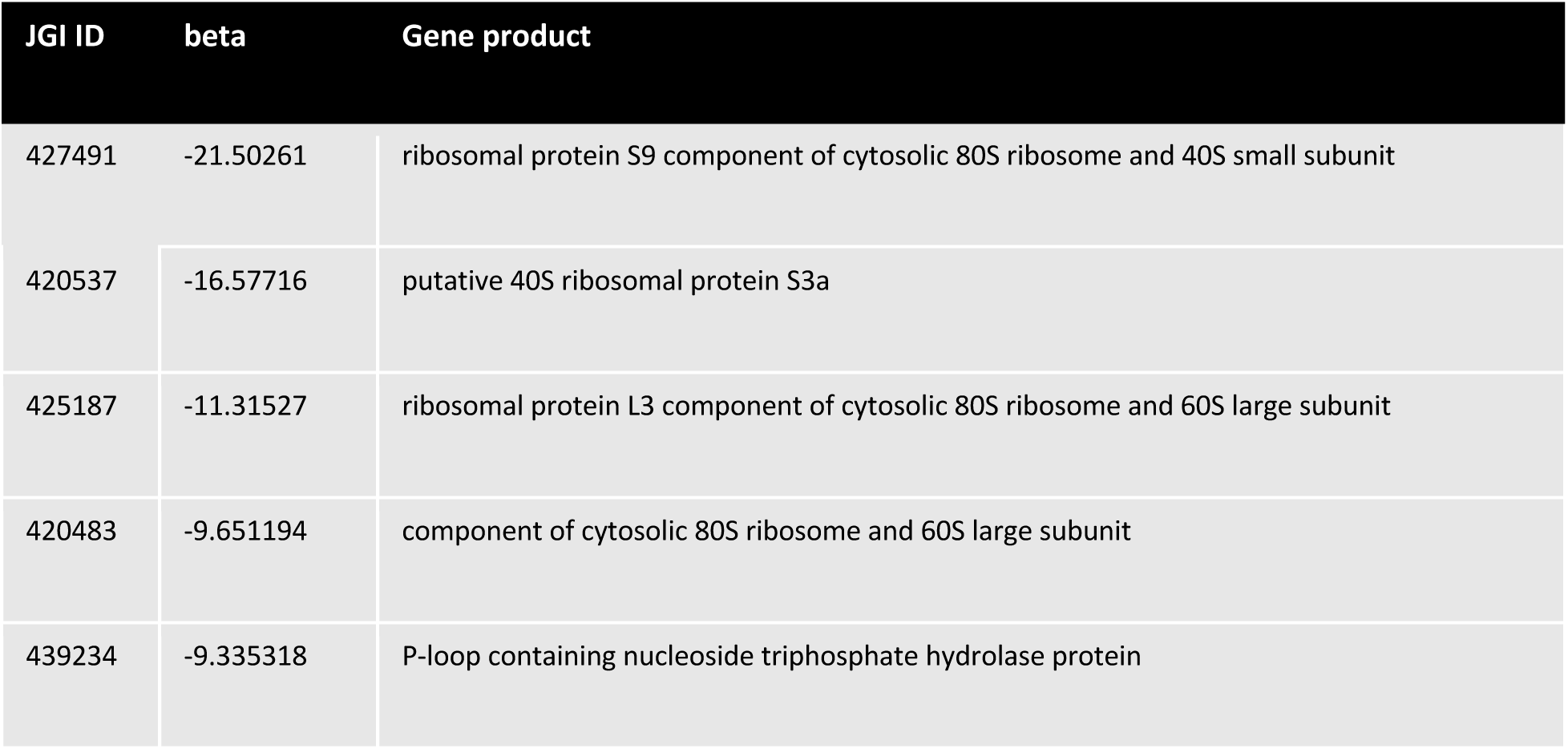
The annotations of the top 5 genes whose expression was downregulated with %DO.

**Table 9:**
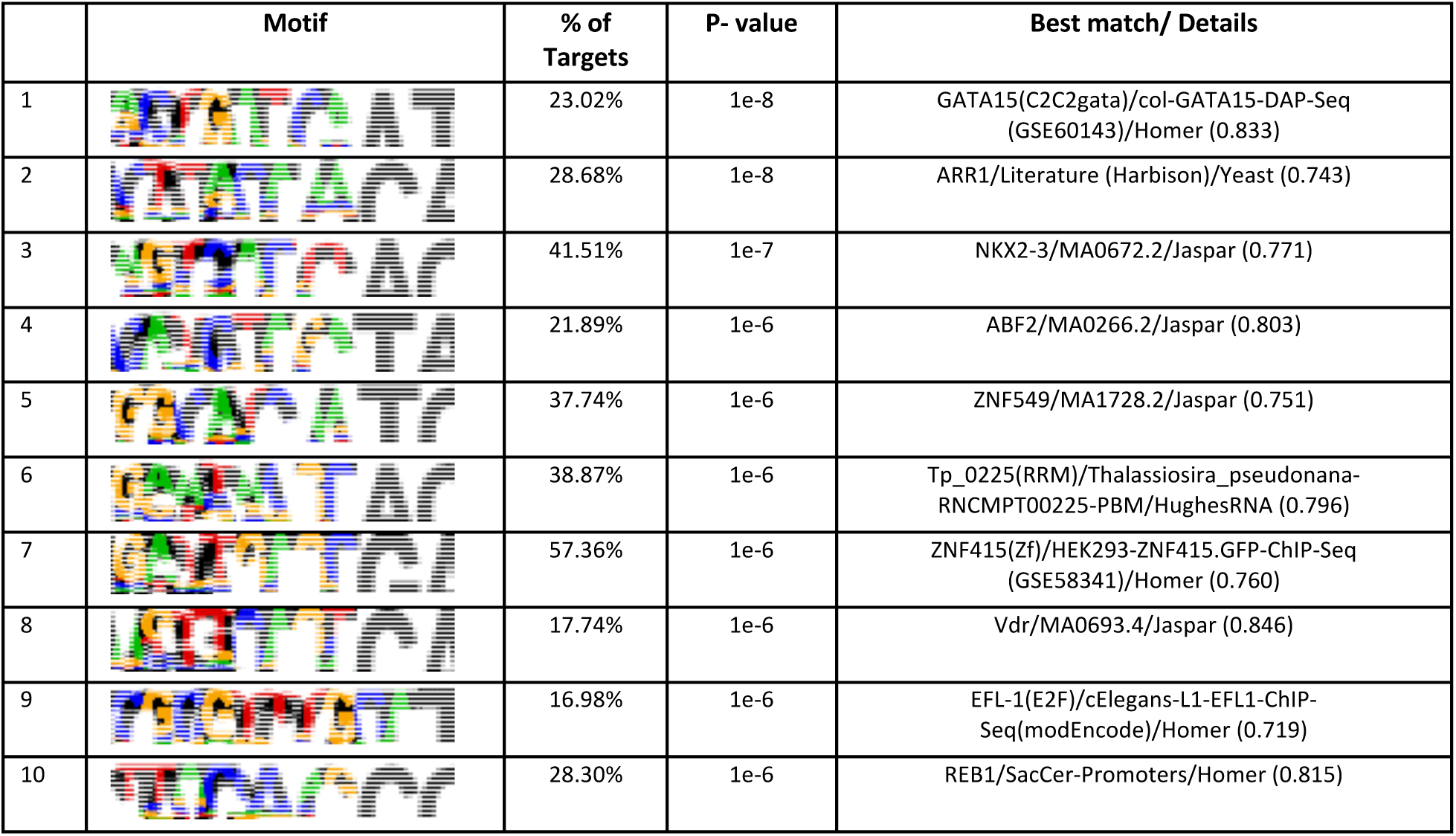
Enriched promoter motifs at the %DO upregulated genes.

|  | Motif | % of Targets | P- value | Best match/ Details |
| --- | --- | --- | --- | --- |
| 1 |  | 23.02% | 1e-8 | GATA15(C2C2gata)/col-GATA15-DAP-Seq (GSE60143)/Homer (0.833) |
| 2 |  | 28.68% | 1e-8 | ARR1/Literature (Harbison)/Yeast (0.743) |
| 3 |  | 41.51% | 1e-7 | NKX2-3/MA0672.2/Jaspar (0.771) |
| 4 |  | 21.89% | 1e-6 | ABF2/MA0266.2/Jaspar (0.803) |
| 5 |  | 37.74% | 1e-6 | ZNF549/MA1728.2/Jaspar (0.751) |
| 6 |  | 38.87% | 1e-6 | Tp_0225(RRM)/Thalassiosira_pseudonana-RNCMPT00225-PBM/HughesRNA (0.796) |
| 7 |  | 57.36% | 1e-6 | ZNF415(Zf)/HEK293-ZNF415.GFP-ChIP-Seq (GSE58341)/Homer (0.760) |
| 8 |  | 17.74% | 1e-6 | Vdr/MA0693.4/Jaspar (0.846) |
| 9 |  | 16.98% | 1e-6 | EFL-1(E2F)/cElegans-L1-EFL1-ChIP-Seq(modEncode)/Homer (0.719) |
| 10 |  | 28.30% | 1e-6 | REB1/SacCer-Promoters/Homer (0.815) |

**Table 10:**
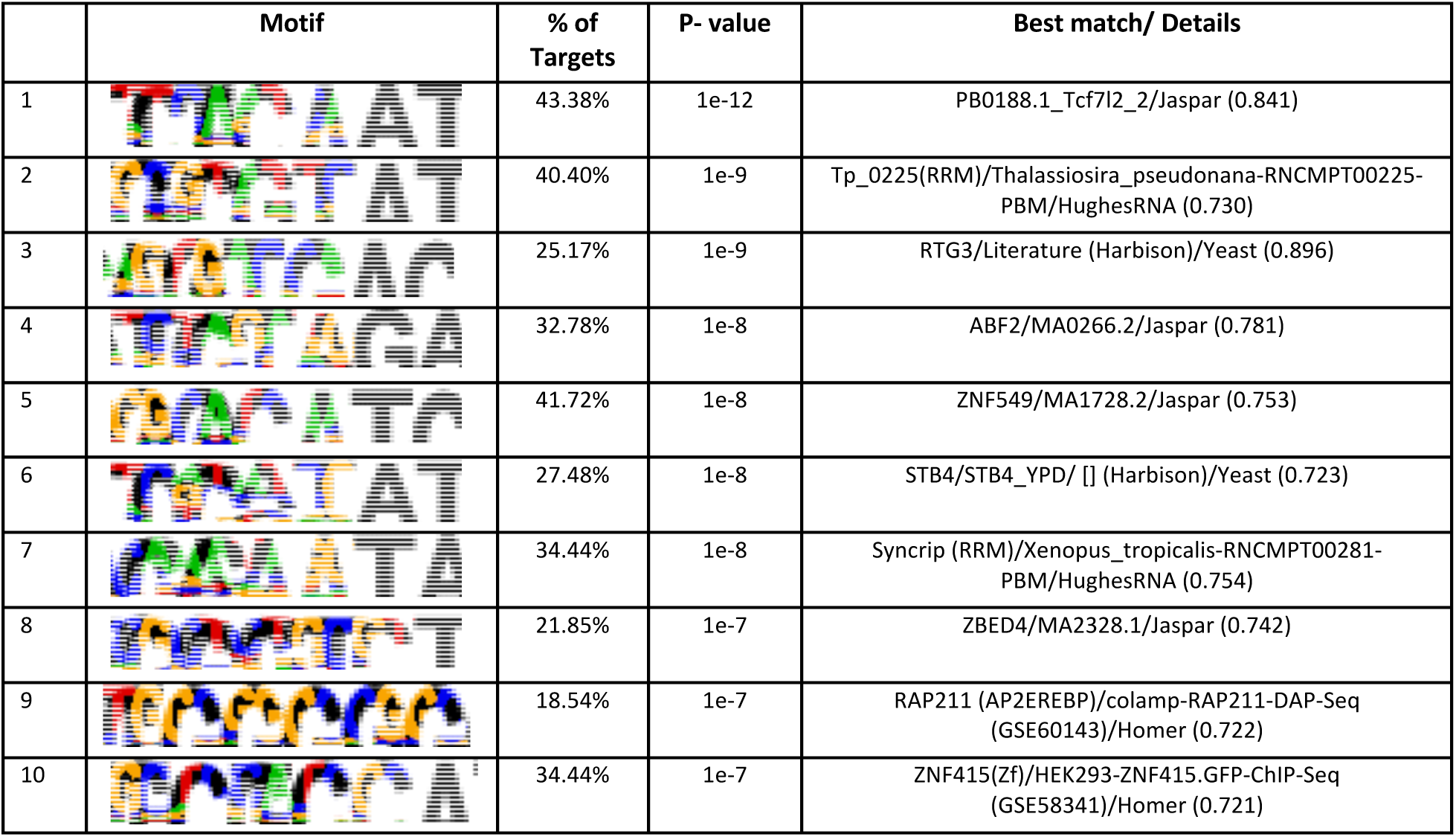
Enriched promoter motifs at the %DO downregulated genes.

|  | Motif | % of Targets | P- value | Best match/ Details |
| --- | --- | --- | --- | --- |
| 1 |  | 43.38% | 1e-12 | PB0188.1_Tcf7l2_2/Jaspar (0.841) |
| 2 |  | 40.40% | 1e-9 | Tp_0225(RRM)/Thalassiosira_pseudonana-RNCMPT00225-PBM/HughesRNA (0.730) |
| 3 |  | 25.17% | 1e-9 | RTG3/Literature (Harbison)/Yeast (0.896) |
| 4 |  | 32.78% | 1e-8 | ABF2/MA0266.2/Jaspar (0.781) |
| 5 |  | 41.72% | 1e-8 | ZNF549/MA1728.2/Jaspar (0.753) |
| 6 |  | 27.48% | 1e-8 | STB4/STB4_YPD/ [] (Harbison)/Yeast (0.723) |
| 7 |  | 34.44% | 1e-8 | Syncrip (RRM)/Xenopus_tropicalis-RNCMPT00281-PBM/HughesRNA (0.754) |
| 8 |  | 21.85% | 1e-7 | ZBED4/MA2328.1/Jaspar (0.742) |
| 9 |  | 18.54% | 1e-7 | RAP211 (AP2EREBP)/colamp-RAP211-DAP-Seq (GSE60143)/Homer (0.722) |
| 10 |  | 34.44% | 1e-7 | ZNF415(Zf)/HEK293-ZNF415.GFP-ChIP-Seq (GSE58341)/Homer (0.721) |

### 3.4 Proteome composition shifts during algal growth and stress response

To gain additional molecular information from the algae as they sensed transitions in their environment, a comprehensive proteomic analysis was conducted on the two pond populations longitudinally. Similar to the transcriptomics data, PCA of the proteomics data (Additional File 4) revealed a highly dynamic phase during the first two days of the culture (Fig. 8). After entering the log phase, however, the proteome becomes progressively more stable, while at the same time maintaining a distinct profile between the pre- and postinfection periods. This was in contrast to the more dynamic changes in the transcriptome after infection, highlighted in the plasticity of the transcriptional regulation in relation to the environmental triggers in comparison with the stability of the protein expression. Further analysis of the correlations between specific transcripts and proteins will be discussed later in this article.

**Fig. 8:**
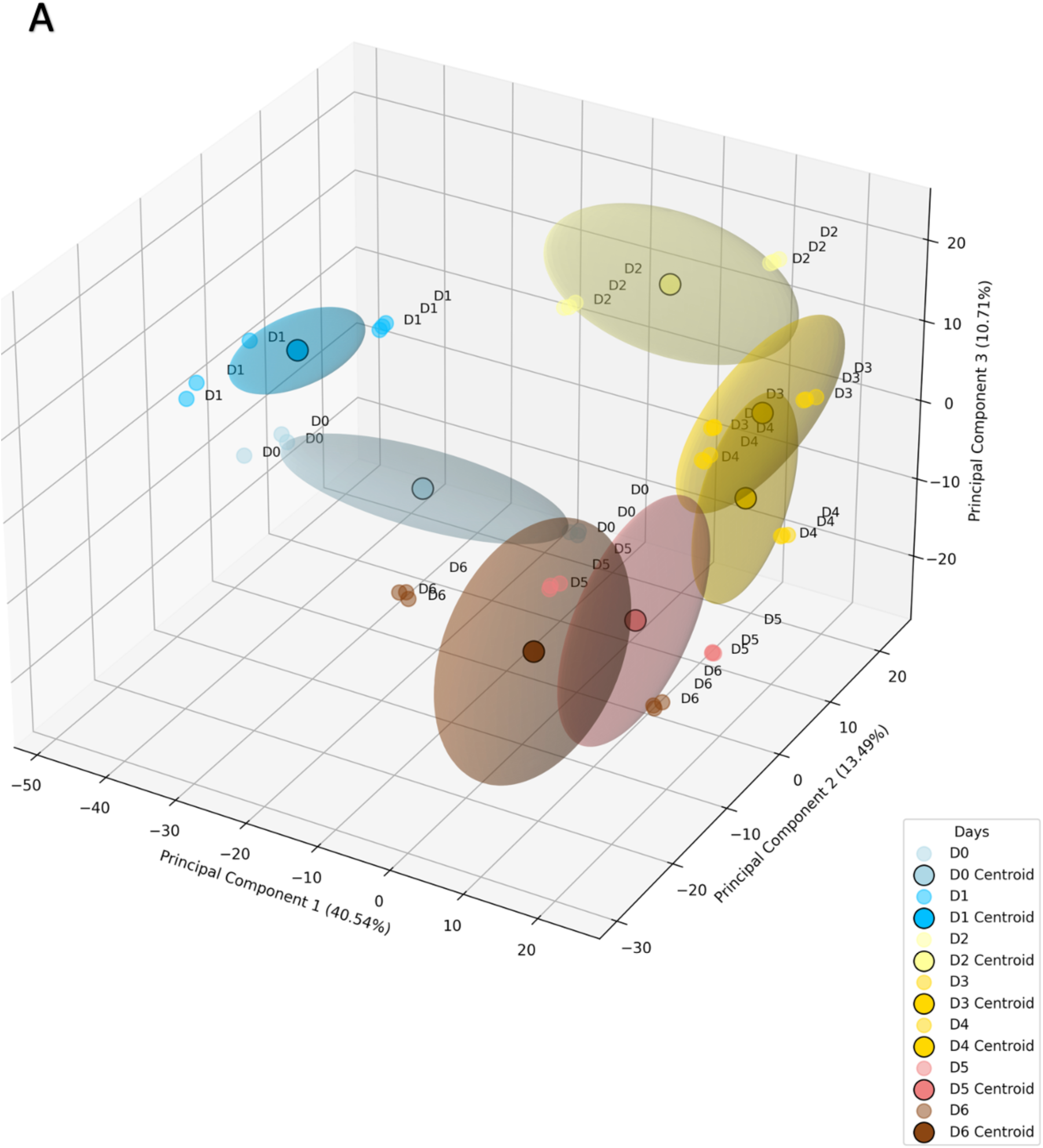
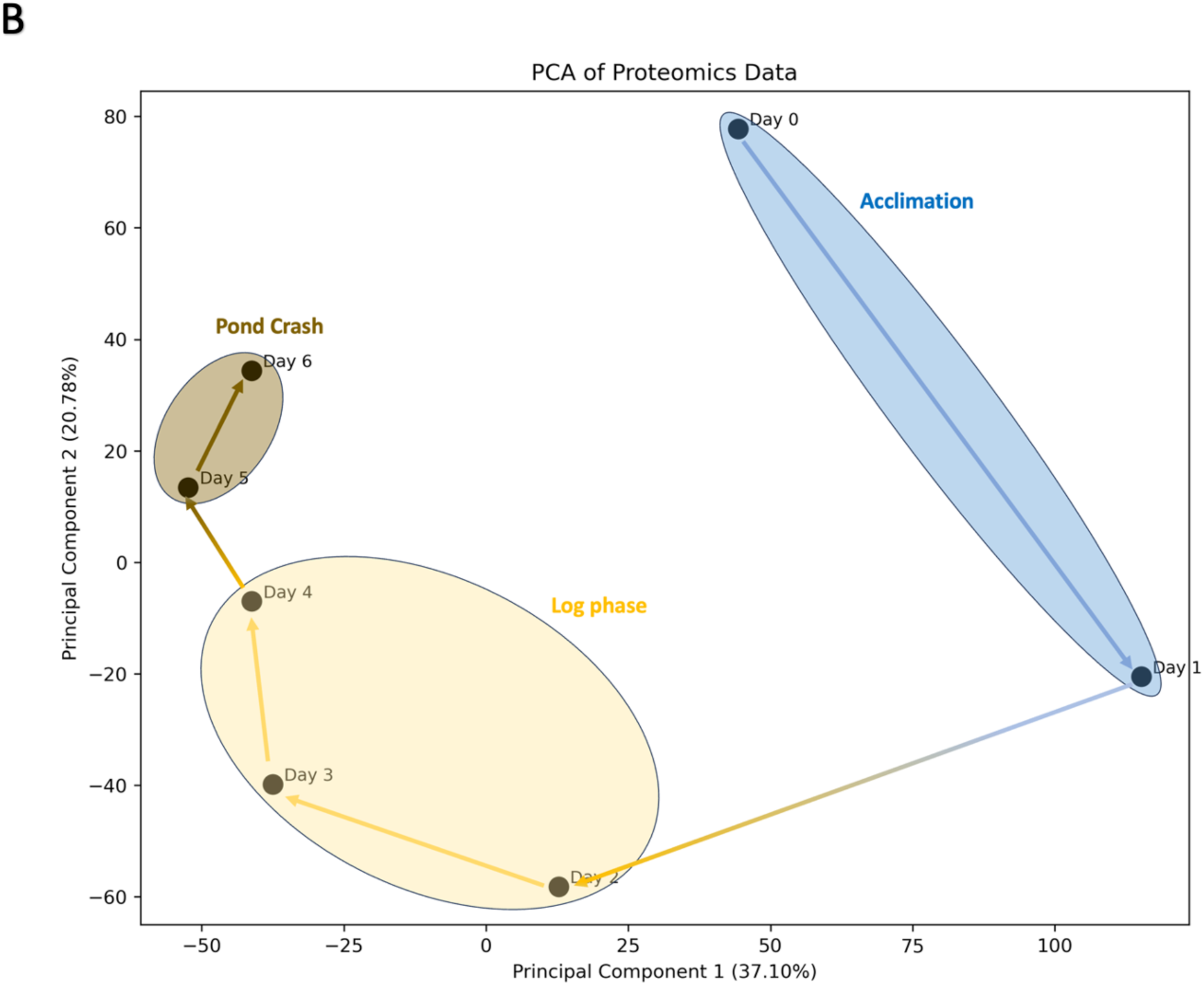
PCA of the proteomics. A) 3D PCA analysis of the proteomics, showing the distribution of all the technical replicates for each day from the two ponds. Centroids are indicated with a darker point, and standard deviation is indicated by the shaded circle. B) 2D representation of the PCA showing the average points for each day. The arrows indicate the trajectory of the time course.

### 3.5 Metabolomic analysis reveals potential indicators of the stress response and pathogen infection

PCA of the metabolites (Additional File 5) revealed distinct profiles during the three phases of cultivation similar to the transcriptomic and proteomic PCAs (Fig. 9A and 9B). Interestingly, day 3, which is in the middle of the log phase, stood out in the PCA. This contrasted with the observation made from transcriptomics. While log phase was linked with transcriptomic stability, the metabolic profile is very dynamic during the growth phase.

**Fig. 9:**
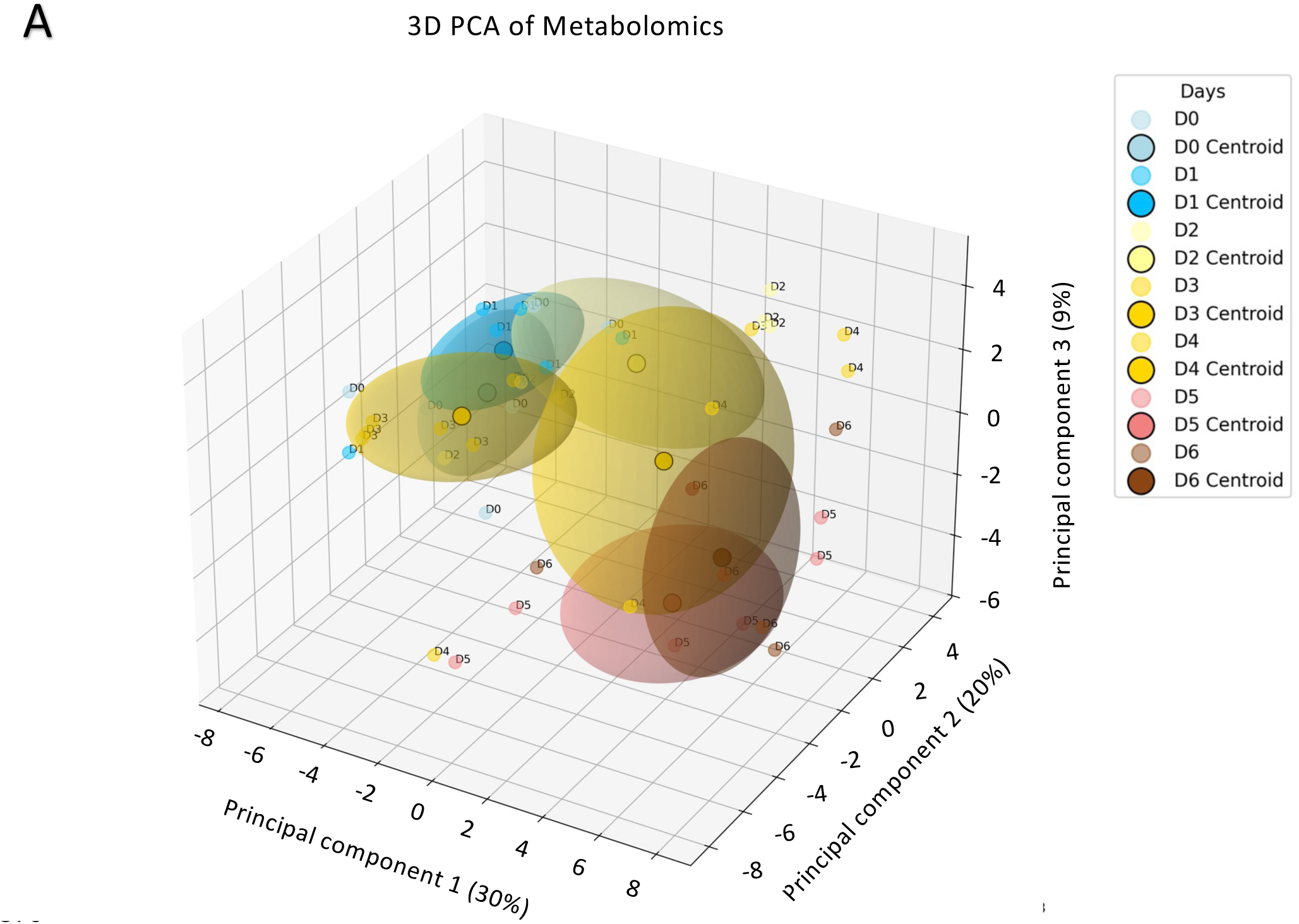
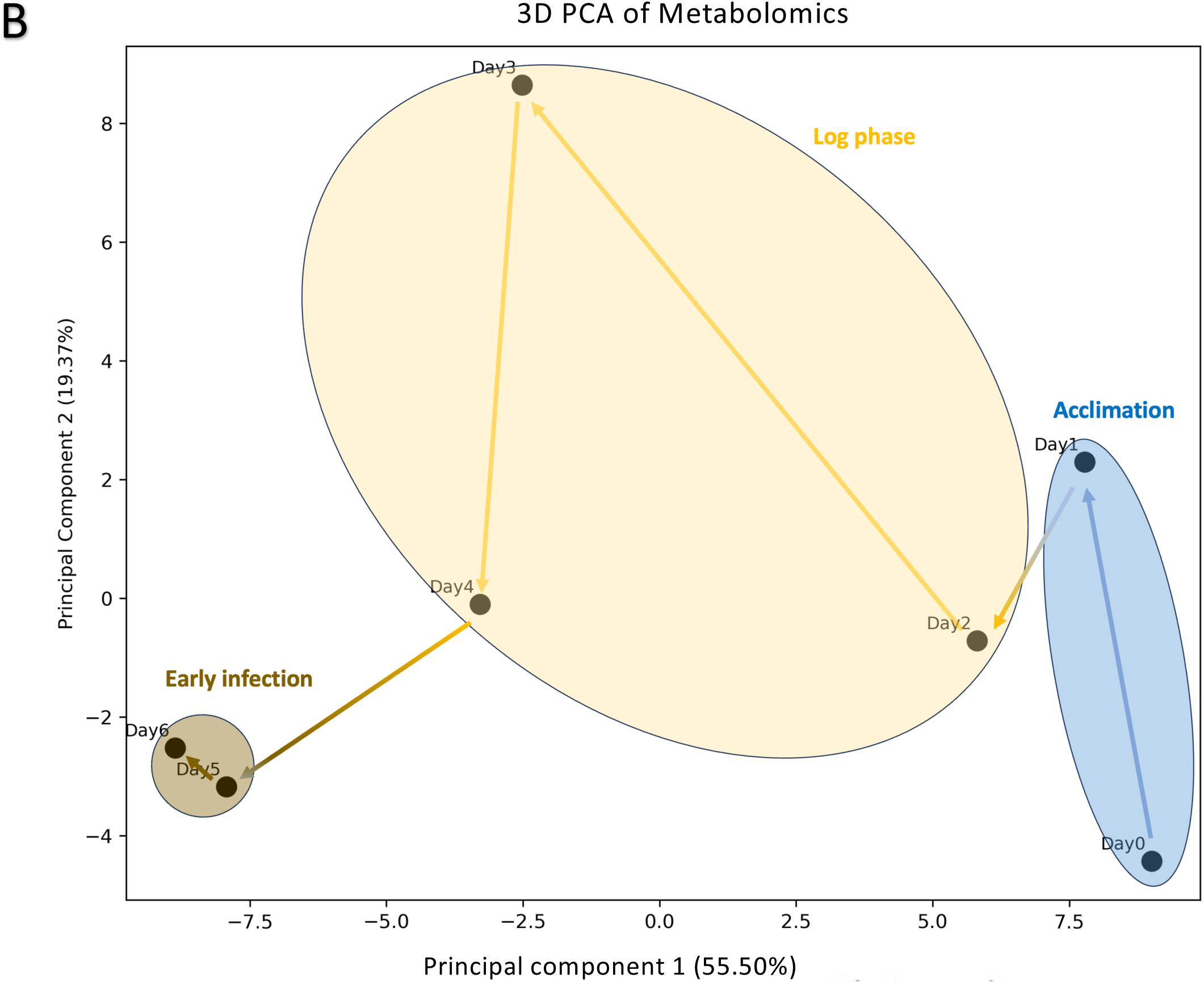
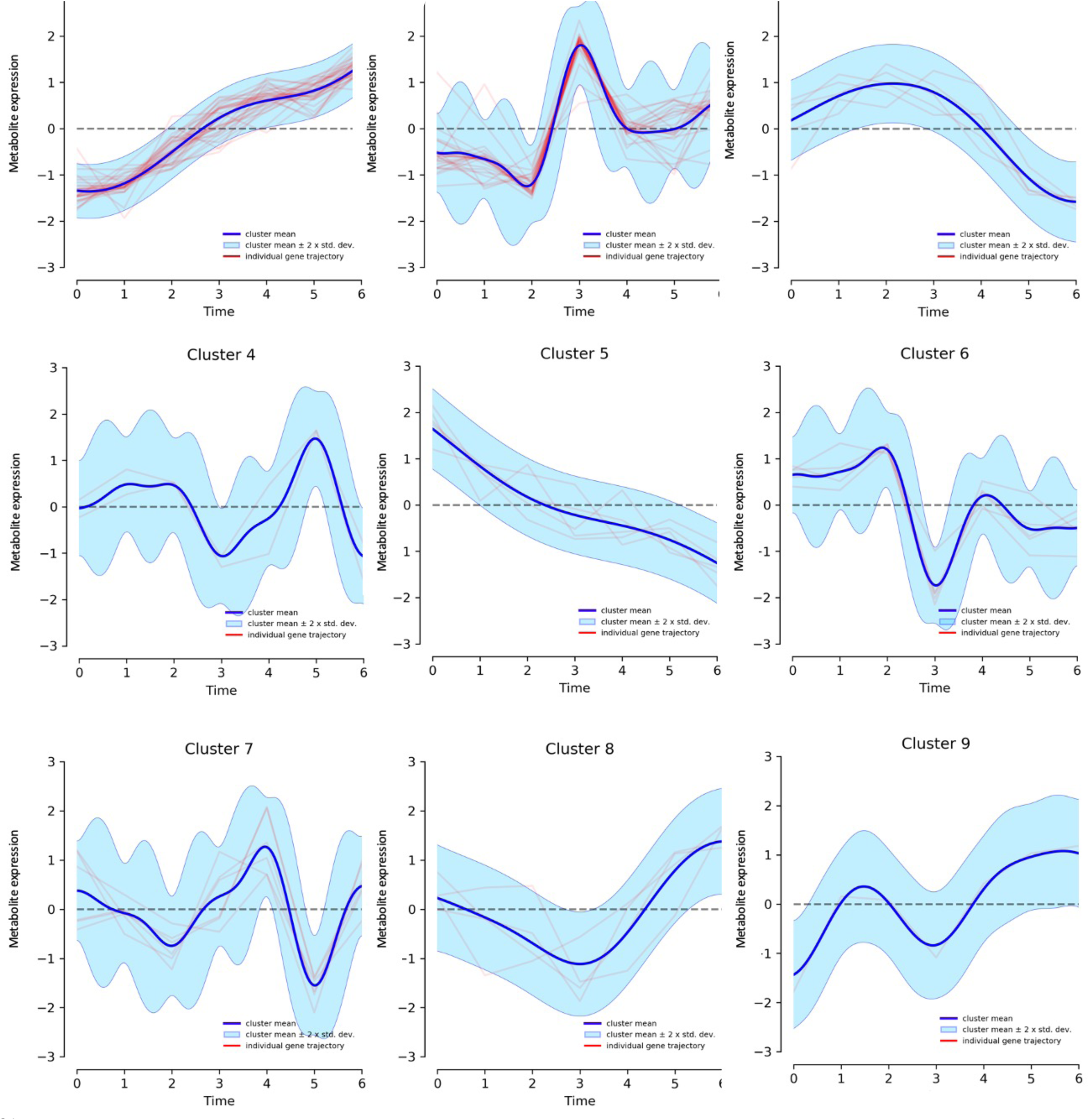
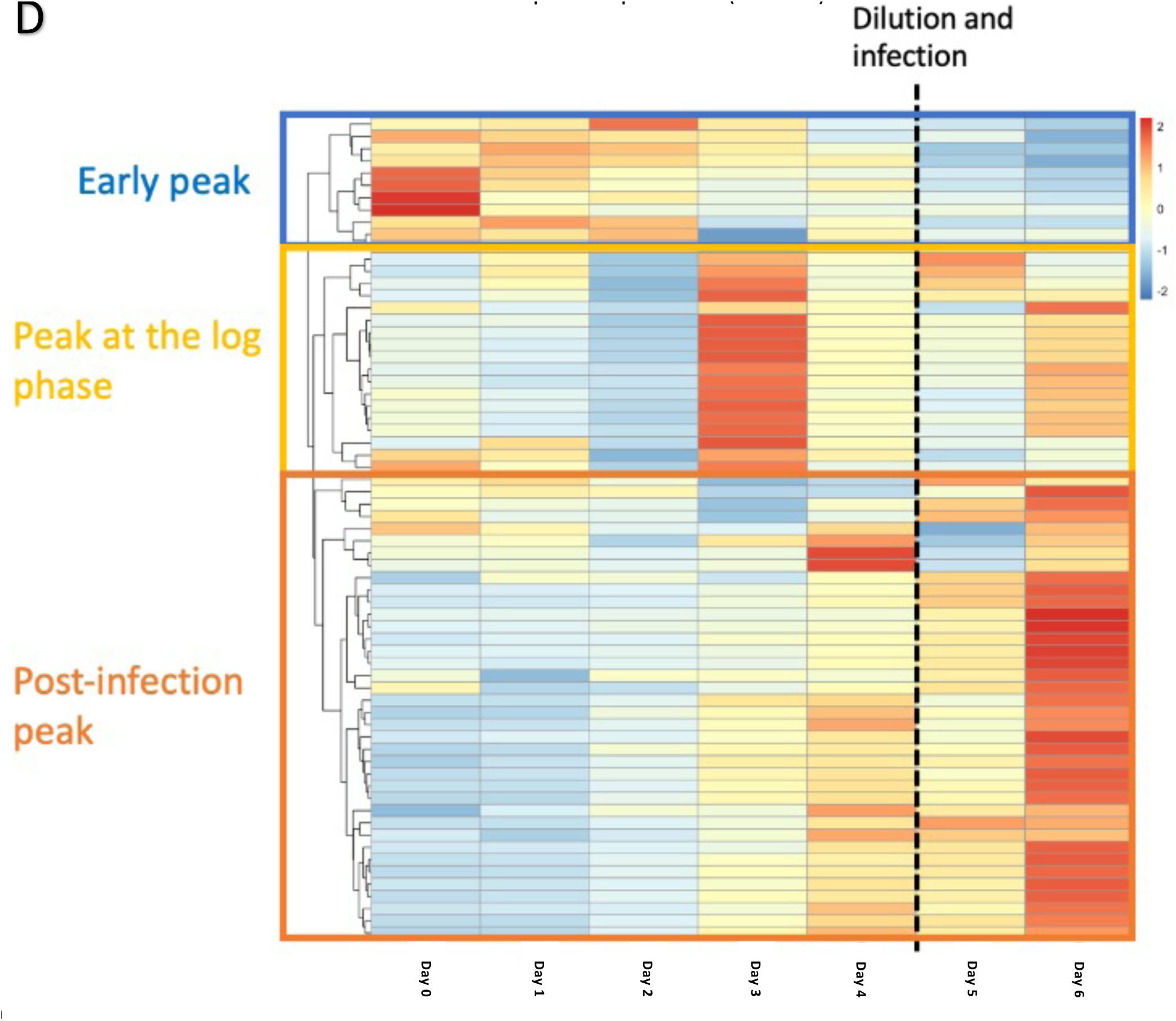
The metabolomic analysis from the pond cultures. A) 3D PCA analysis of the metabolomics, showing the distribution of all the technical replicates for each day from the two ponds. Centroids are indicated with a darker point, and standard deviation is indicated by the shaded circle. B) 2D representation of the PCA showing the average points for each day. The arrows indicate the trajectory of the time course. C) Clustering analysis of the abundance of the metabolites throughout the studied period, via DPGP. D) Heatmap with the hierarchical clustering of the significantly produced metabolites throughout the run of the cultivation. It includes the metabolites which had more that 10x higher signal than the metabolites in the blank.

We performed clustering analysis of the longitudinal metabolomics data to further understand the metabolites that covaried (Fig. 9C). We used DPGP [44], a nonparametric clustering algorithm that can factor in longitudinal information, to separate metabolites into clusters on the basis of their patterns in abundance. We found that many metabolites either had their peak or trough on days 3 or 4, the midpoint of the log phase (clusters 2, 4, 6 and 8 and 9). The most abundant cluster (Cluster 1) included metabolites whose abundance steadily increased over time. On the other hand, Cluster 5 showed the opposite trend, starting with high abundance and slowly decreasing in abundance over time.

Hierarchical clustering of the scaled and centered metabolite abundances in the heatmap, shown in Fig. 9D, highlights the emergence of three groupings of metabolites, and is in relative accordance with the DPGP clustering. The first group consists of the metabolites whose abundance peaks in the lag phase. The second group peaked in the log phase. The last group, which was the largest by number, contained the metabolites that reached their peak abundance after the infection. The last group included hormones and steroids, such as 5-Dihydrotestosterone, 7-Hydroxytestosterone, Estrone, Estriol and 19-Norandrostenedion.

To further verify our findings from the metabolomic analysis on the ponds, specifically on the discovery of infection biomarkers, we repeated the experiment at the flask level in laboratory settings (Additional File 6). The cultures were infected with the same amoeboaphelid between day 4 and day 5 and samples for metabolomics were taken on specific days and analyzed. It should be noted here that the smal-scale experiemnt was conducted in a closed and axenic environment, unlike the large-scale experiment, meaning that the microbiome composition difference could account for changes in metabolic profiles. However, we were able to confirm some potential compounds that are significantly upregulated in both conditions after the infection, and upon further investigation could serve as potential biomarkers for upcoming culture crashes (Fig 10)..

**Fig. 10:**
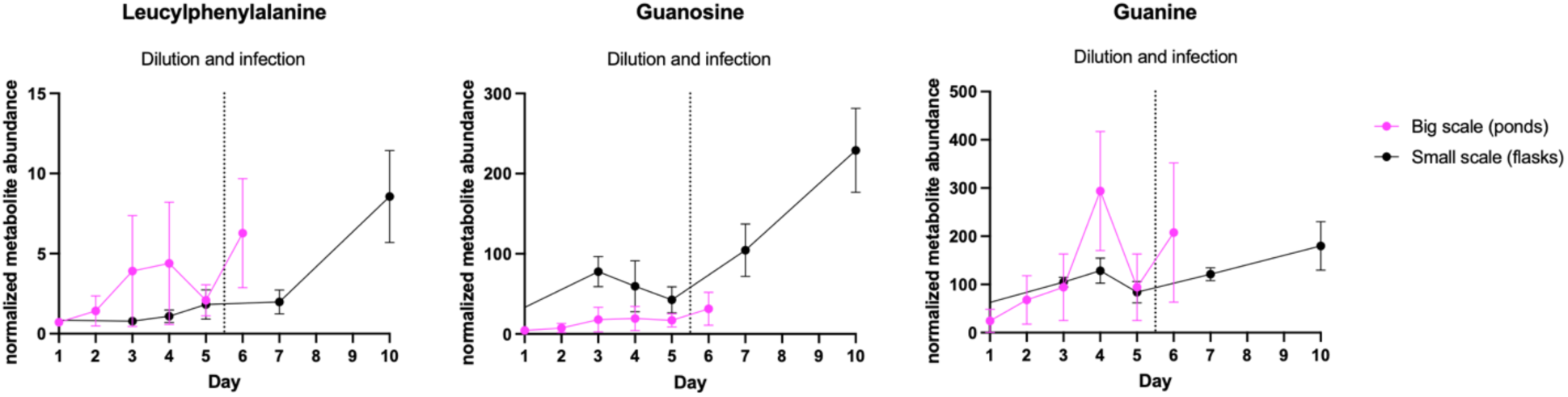
Metabolites who can act as early pond crash biomarkers, verified by small-scale experiment.

### 3.6 Integrated multi-omic analysis revealed distinct molecular features related to scale transition and infection defense

To assess the global relationships among proteins, metabolites and transcripts across our longitudinal experiment, we performed an integrative analysis via the DIABLO framework in the mixOmics R package [38]. We performed this analysis using all omics data from days 0 and 1 to explore the molecular features of the scale-up stress, and separately used days 4 and 5 data to explore that of the infection stress. While we found that there was no overlap in the key proteins or transcripts output by the model between the scale up and infection stress, here were two metabolites, guanine and 12-oxo-Phytodienoic acid, were common across the two stressors, suggesting more convergence at the metabolite level than at the molecular pathway level.

We took the molecular features from the first latent component of both analyses and combined them into one heatmap to comprehensively visualize these results (Fig. 11). As expected, the columns clustered by day, and we observed that the two transitions had almost entirely unique patterns. We observed a cluster of transcripts (all significantly differentially expressed between days 0 and 1), proteins, and myristic acid elevated on day 0 relative to the other days. Similarly, we observed another distinct cluster of transcripts and proteins, as well as 1,5-hexadiene and 2,3-Diaminosalicylic acid, which were elevated on Day 4 compared with the other days. However, a decrease in a different cluster of proteins, as well as maltose and hypogeic acid, distinguished day 1 relative to the other days. These results suggest distinct molecular and metabolic profiles in response to different stressors, identifying potential unique biomarkers and target pathways for detection and intervention.

**Fig. 11:**
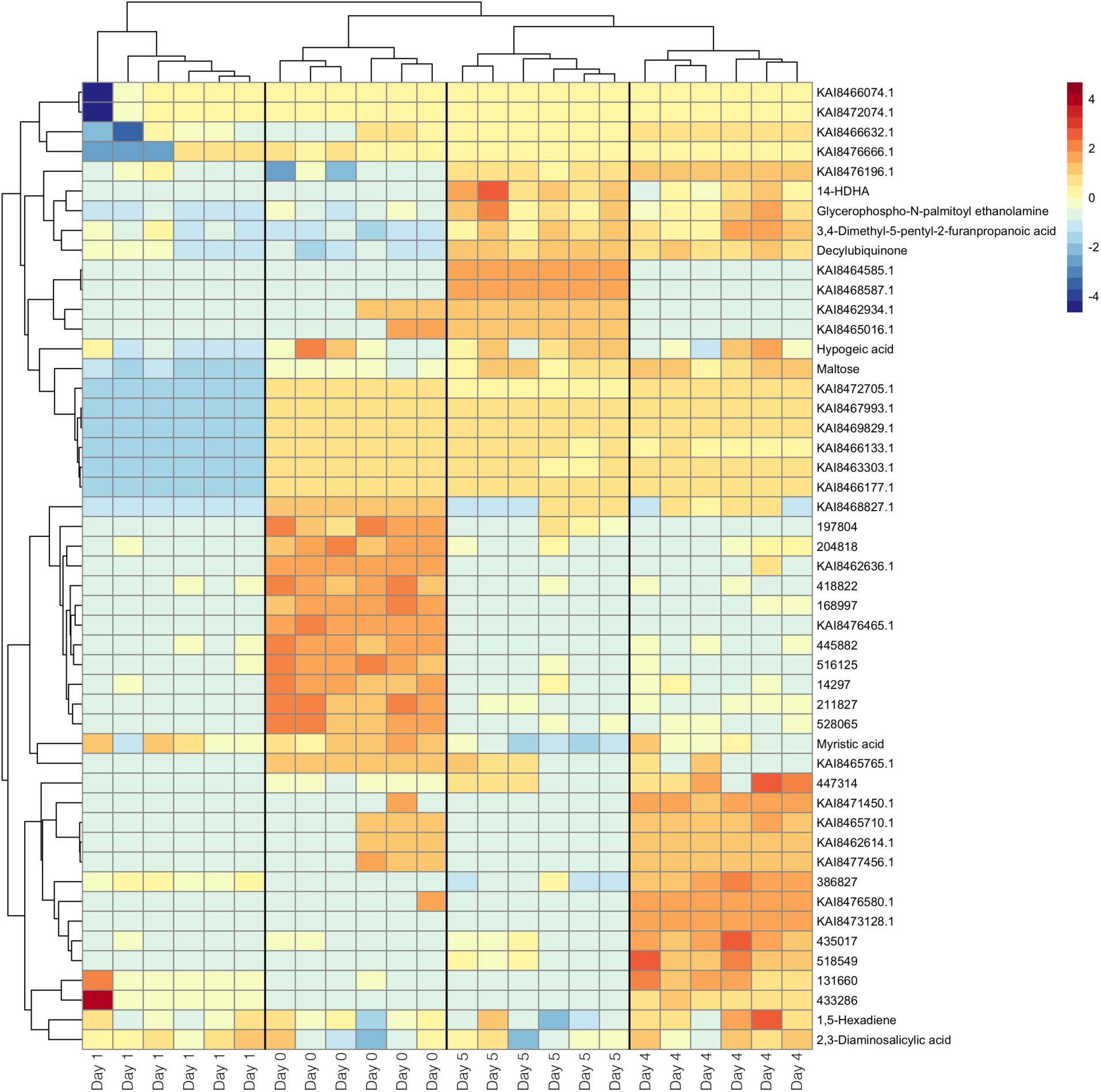
Indicative signatures of scale adaptation and infection. Heatmap containing the molecular features (genes, proteins and metabolites) which change significantly their abundance during the transition from day 0 to day 1 and/or from day 4 to day 5. Their abundance is being indicated by a scaled blue-to-red scheme. Each of the 6 biological replicates for each condition are indicated separately

In order to determine the dynamics of the transcriptomic and proteomic observations for pathway analysis we performed a correlation analysis among the daily transcriptomic and proteomic datasets and between the two datasets (Fig.12A). The results indicate a well-expected divergence in the two datasets between the first days (corresponding to acclimation regulation) and the rest of the growth. They also showcased that the proteomics are more stable and less dynamic than the transcriptomics, and thus less sensitive as observation markers for stress events.

To integrate the various data streams and confirm the potential diverging signals and pathways across the two stress events, we wanted to demonstrate that specific genes that shared a regulatory motif partnered with proteins and metabolites to regulate distinct pathways in a stressor-specific manner. We performed a protein-protein interaction network analysis to showcase the specific networks which regulate the scale-up and the infection stresses, via STRING (Fig.12B). While common features exist in both networks, it is evident that distinct and unique interactions are significant between those two stressors, verifying our initial observations. The enlarged network figures are shown in Additional Figures 2 and 3.

Specifically, we see nodes like Glutamate Dehydrogenase 2 (GDH2), Malate Dehydrogenase (MDH) and the bacterial-type polymerase AGS1 that are shared between stressors, and these genes are known to be involved in processes such as carbon and nitrogen fixation, and macromolecule biosynthesis [45]. We also see families such as Epidermal Growth Factors (EGFs) [46], the ribosomal small protein subunits (RPSs), Phosphoribosylpyrophosphate Synthetases (PRPSs) [47] and the genes for prolactin (PRPLs) representing the larger gene expression and translational machinery. These are common nodes between the stressors, but we see different members of these families in the two stressors, suggesting that even pathways that might be shared between stressors are likely controlled through different sub-pathways depending on the nature of the stressor. HSPs (heat shock proteins) seem to be specific to the scale-up network, which is consistent to needing to keep proteins properly folded during rapid new protein synthesis and cell cycle progression during scale up [48] [49]. With amoeboaphelid infection, ICL1 (isocitrate lyase), a player in the glyoxylate cycle, appears to be a unique hub, which would be consistent with re-directing metabolic processes during stress as has previously been shown in other species [50] [51]. This suggests that there are shared factors, shared processes with distinct factors, and distinct processes all at play for stressor-specific responses, and within these pathways could lie biomarkers and countermeasure targets that are either stress-specific or stress-agnostic.

## 4. Discussion

### 4.1 Transcriptomics reveal the commonalities and differences between the molecular mechanisms governing scale-up stress and infection response

We performed a multi-omic analysis of *M. minutum* 26B-AM cultivated over 12 days in 1000 L open raceway ponds to explore the molecular signatures that manifest in response to stressors often encountered at a large scale. The three distinct growth phases we observed in the 1000 L ponds were in line with what is typically observed at a large scale, i.e., batch cultivations of algae [52] [53]. Culture scale-up is often evidenced by a lag phase, as we observed in our cultures, especially when moving from well-controlled laboratory cultures into open raceway ponds [52] [54] [55]. A change in scale represents a significant stressor for the cultures, introducing a combination of different environmental signals, including changes in light, radiation, heat, gas distribution, as well as microbial and environmental particle composition that requires an acclimation period, which is reflected in our growth curves over days 0 and 1.

Similarly, algal pests are common stressors experienced in open raceway ponds and active areas of study. Changes in phenotypic measurements in response to infection or even visual indications of infection stress, typically appear sometime after the initial infection. In our cultures, we introduced the amoebaphelid on day 4 but did not begin to observe a decline in growth, which was emblematic of a culture crash, until day 8, accompanied by a color change in the culture from light green to brown. Pond crashes, which are driven by algal pathogens and often damage crops overnight, account for almost 30% of algal crop losses annually [56], increasing the overall cost of algal applications. Despite the dramatic effects of those infections on ponds, the cellular mechanism of the infection which includes the insertion of the parasite in the host cell [19], allows a certain amount of time of incubation before a noticeable crash [57]. Cellular and molecular alterations are already in place and if they can be detected in a timely manner, the irreversible effects of a pond crash can potentially be avoided, or at least resource allocation decisions can be made much earlier.

Differential expression analysis revealed that culture scale up has a substantial effect on the transcriptome, as evidenced by the much larger portion of significantly differentially expressed transcripts between days 0 and 1 compared with the other days, including those impacted by infection. In addition, infection is not expected to trigger a wide immune response from the algal cells, reflected by transcriptional adaptation. Characterizing these pathways and adjusting conditions to prevent their activation could help acclimate the culture in a less stressful way and potentially make it more resilient to future stressors.

With respect to infection, current strategies for early indication of contamination include visual detection strategies such as flow cytometry [58], in situ microscopy [59], multispectral image analysis [60] and photosynthetic measurements [61] [62]. Those methodologies, although relatively quick, require a certain pathogen titer in the culture. Biomolecular approaches include 16S and 18S sequencing [63], and the detection of Volatile Organic Compounds (VOC), as indirect early indicators of an infection [64]. Here we show that the algae can also be used as early indicators of infection, as demonstrated by the changes in transcripts.

Because many of the transcripts in *M. minutum* are poorly annotated, we used motif analysis to further explore the significantly differentially expressed genes. Most identified motifs were unique to each transition, potentially indicating two distinct regulatory pathways between the scale adaptation and pathogen stress. However, common elements between the two datasets exist, implying that there are factors that could be used to coregulate responses to different stressors (either endogenously or via synthetic biology). An example of a potential point of convergence are the motifs recognized by the transcription factors mitochondrial retrograde 1 (RTG1) and 3 (RTG3), both of which were originally identified in yeast and are involved in mitochondrial metabolic regulation.

Among the promoters of genes differentially expressed during scaling up, we also identified a motif labeled by HOMER as ZNF549, which is likely bound by zinc finger proteins, a family of transcription factors that have been shown to regulate stress responses in other algal species [65] [66]. In addition, 31.84% of the motifs correspond to a promoter motif from the diatom *Thalassiosira pseudonana* suggesting possible functional importance across diverse algal species. A notably high percentage (66.66%) of the promoter regions of the genes that were significantly differentially expressed after the infection contained a motif that can be recognized by Auxin Response Factor 1 (ARF1), a transcription factor identified in *Arabidopsis thaliana.* ARF1 is responsive to the phytohormone auxin, which is responsible for a wide range of developmental and stress responses in plants [67] [68]. Given the evolutionary proximity between higher plants and green algae, it is likely that ARF1 may have similar functions in algal responses post-infection and might suggest a possible defense mechanism. The identification of these factors points to potential targets for synthetic biology in *M. minutum* in the future, to either boost or tamper with specific molecular responses.

### 4.2 Metabolomics provide useful information about potential stress and growth biomarkers in algae cultures

A survey of the metabolites released by the algae upon encountering various stressors also provides valuable insights into predictive markers, pond health indicators, and infection indicators, while also explaining the natural mechanisms of growth and defense in algae. We used PCA and DPGP to look at global trends in metabolites over time. While log phase was linked with transcriptomic stability, the metabolic profile is very dynamic during the growth phase.

These metabolites are highly important, both as early and timely indicators of an upcoming crash and as guides for the development of prophylactic strategies for algal cultures. Among those metabolites, [69]there are multiple hormones and steroids (5-Dihydrotestosterone, 7-Hydroxytestosterone, Estrone, Estriol and 19-Norandrostenedion). While such compounds originate in the aquatic environments from sources other than algae, many algal species are known to biotransform them or completely remove them from the system [70] [71] [72], with effects that can be either toxic or growth promoting for the cells [73] [74].

As the ponds were open to environmental influence, bacterial communities present are composed of both environmental bacteria and the algae-associated microbiomes living on the surface of the algal cells [75]. Although *M. minutum* 26B-AM has been extensively studied as a potential source of biofuels [52] [76] [77] [78], there is only one other publication related to its associated microbiome during cultivation for bioproducts [79]. It was described there that the bacterial community was dominated by Alphaproteobacteria. No information was available from this study on eukaryotic community members other than *Monoraphidium*, although since cultures were grown in outdoor raceway ponds similar to the ones used in the current study, it can be assumed that there were both prokaryotic and eukaryotic environmental members. Other studies of related green algae species found that bacteria might play important roles in regulating nutrient dynamics and impacting cross-species interaction between eukaryotic and prokaryotic community members [80] [81]. There was no statistically significant relationship between any of the bacterial phyla and algal productivity, suggesting that the aphelid infection was the primary driver of lowered algal productivity during the infection period. However, since the aphelid was not detectable until seven days post-infection, this suggests that early pest detection should rely on both molecular methods and microscopy, which may allow for earlier detection and thus a longer response time for algal cultivators prior to crop loss.

We also identified 14-HDHA, a hydroxy docosahexaenoic acid oxylipin which is derived from docosahexaenoic acid (DHA). The production of oxylipins from algae is linked with oxidative stress [54], suggesting a similar regulatory pathway here for pathogen defense. When we repeated the infection treatment on the flask level, we observed similar uptake on DHA post-infection. DHA is a type of omega-3 fatty acid with great commercial interest of human nutrition, often produced by algal systems [82] [83] [84] [85]. It has been also reported that co-cultures of algae and fungal species can improve the yield of algal-produced DHA [86] [87].

The comparison of the metabolomics from the ponds and the flasks pre- and post-infection gave us three metabolites that increase after the addition of the pathogen in both scales. Those were Leucylphenylalanine, guanosine and guanine (Fig. 10). While the comparison between a big-scale open system and controlled laboratory axenic environment can be confounded by environmental and experimental factors, the verification of the uptake of those metabolites certifies their status as potential biomarkers for upcoming culture crashes.

### 4.3 Transcriptomic and phenotypical observations reflect light and gas-regulated responses in M. minutum

Light and gas constitute the two cornerstone environmental factors for aquatic organisms, especially the photosynthetic organisms such as green microalgae. Indeed, light and CO_2_ availability are essential not only for photosynthesis, the primary metabolic function of microalgae but also for photoprotection, lipid composition, and carbon concentration [88] [89] [90] [91] [92]. The conserved and well-studied role of light and CO_2_ availability led us to explore which *M. minutum* genes are corelated with these conditions, to test the validity of our method on identifying molecular signatures. We observed that translation-related proteins were strongly correlated with high light intensity. Protein translation is an essential regulatory step for phenotypic acclimation, and its control by light has been observed in other algae, including the Chlorophyte *Ulva prolifera* [93], the Charophyte *Klebsormidium nitens* [94], and *Chlorella sp.* [95]. In addition, we noted a gene coding for a structural component of Photosystem I among the genes whose expression was downregulated in response to light. Photosystem I is a crucial component of the electron transport chain and is responsible for the light-mediated transfer of electrons across the thylakoid membrane. Excess light can cause oxidative stress and subsequent photoinhibition in Photosystem I, and in many photosynthetic organisms, including in *Chlamydomonas reinhardtii*, light can suppress the expression of structural proteins in Photosystem I to protect the machinery from this type of damage [96]. Future work to test this hypothesis across algal species will be critical.

Motif analysis of the promoter regions of genes correlated with light highlighted two motifs related to homeobox proteins in higher plants. The *Zea mays* homeobox 2a (ZmHOX2a) protein, whose expression in maize is restricted to meristems, is involved in in plant development and growth [97]. In addition, this motif might more broadly represent homeobox transcription factors, which are known to regulate environmental responses in plants [98] [99], suggesting evolutionary conservation of light-induced gene regulation through homeobox transcription factors across the plants and algae. A motif corresponding to the same transcription factor is also found among the motifs from the promoters of the downregulated genes by light, highlighting the involvement of these motifs in both the up- and downregulation responses to light. Interestingly, 30.80% of the targets of the promoter regions from the up-regulated genes contained a motif corresponding to the transcription factor RGT1 (Restores Glucose Transport) from *Saccharomyces cerevisiae*. The protein regulates glucose transport though the expression of glucose transporters [100], and the connection between light-induced stress and glucose transport has been documented in other organisms [101] [102]. The motifs of the downregulated genes include one that is recognized by the *Saccharomyces cerevisiae* stress-related multicopy suppressor of SNF1 mutation protein 4 (Msn4), which controls the metabolic response under cellular stress conditions [103].

Among the genes correlated with DO, multiple genes are related to photosynthesis and carbon fixation. A phosphoribulokinase/uridine kinase, an enzyme responsible for a key step in the Calvin-Benson cycle, and a light-harvesting protein of Photosystem I are among the most upregulated genes. Genes related to programmed cell death were positively correlated with DO, an observation that aligns with results from other green algal species linking ROS signaling and programmed cell death [104] [105] [106] [107] [108]. These observations highlight specific molecular changes experienced by large-scale cultures that are likely directly related to environmental fluctuations. Our results also revealed that increased DO caused more gene downregulation than gene upregulation overall, especially for the genes indicated in Table 8. Most of these genes are correlated with structural components of ribosomes, indicating a strong suppressive relationship between O_2_ levels and translational regulation, as was also observed in the case of genes upregulated with light. In fact, the most downregulated gene from the DO was the most upregulated one from light. Since high light is a stress factor, as is low oxygen, this reciprocal relationship between their regulated genes aligns with expectations. The list of the top five downregulated genes includes a P-loop containing nucleoside triphosphate hydrolase, which, in higher plants such as *Arabidopsis,* acts as a signaling molecule involved in ribosome formation and attenuates ROS formation and oxidative stress[109] [110].

Motif analysis of the promoter regions of all the genes correlated with DO revealed enrichment of the binding site for the same transcription factor from the diatom *Thalassiosira pseudonana* that we observed during scale acclimation. This putative transcription factor motif is found in 40.40% of the promoter regions of the genes differentially expressed during scale acclimation (Table 1), which is a sign of a strong contribution of the DO differences to the scale transition stress. Future efforts to identify this transcription factor that is conserved across algae might be fruitful for understanding stress mechanisms and for developing broad-scale synthetic biology-based interventions. The second most enriched motif among the promoters of the upregulated genes corresponds to the transcription factor Arrestin 1 (ARR1), which in higher plants is responsive to the phytohormones Cytokinins and plays a fundamental role in numerous developmental stages of the plant [111] .The motif bound by the yeast transcription factor Regulation of the Mitochondrial Retrograde 3 (RTG3) is also found at the promoters of downregulated genes. This transcription factor regulates in multiple metabolic pathways, including amino acid and nucleotide synthesis [112] indicating a possible role of oxygen-regulated metabolic responses in *Monoraphidium*. Interestingly, motifs corresponding to the same transcription factor were found among the photometers of the differentially expressed genes during the acclimation period and after the infection with the amoeboapheids (Tables 1 and 2), which implies a role for transcription factors that bind this motif across stressors and environmental changes.

We also noted that we identified overlap in the genes correlated with light and DO and those related to stress responses (specifically responses to cold and salt), as determined by [17]. Specifically, of the 24 genes they outline as being annotated with a fasciclin-like domain, a domain with known roles in algal and plant stress responses, we note 2 with expression coordinated with dissolved oxygen (annotation # 450375, 525465), 1 with expression coordinated with light (annotation # 480939) and 2 with expression levels coordinated with both (annotation # 421681, 446308). These results support the validity of our signature identification through multi-omic data integration in algae, and point towards the need for follow-up on our stress signatures for identification of predictive markers for tracking large-scale pond culture resiliency, and molecules for therapeutic intervention.

## 5. Conclusions

In this study, we used multi-omics to investigate the dynamic responses of the green alga *M. minutum* to various environmental changes and stressors. A study of multi-omic analyses of *M. minutum* was recently published, providing a molecular profile of this industrially significant species under high salt and cold temperature stress [17]. We utilized a multi-omic approach in larger open and, thus, more heterogeneous culture systems in order to provide a roadmap for the application of molecular observations at the cellular level and data processing to address the significant challenges faced by the large-scale bioprocessing industry. [57, 113]Our approach aims to address these issues while gaining information about the cellular responses of the species. We demonstrated the ability to integrate datasets of genes, proteins and metabolites that respond to specific stressors or correspond to certain culture stages and transitions, opening up possibilities for follow-up studies in the fields of strain engineering, biomarker development, bioprocessing, and basic cellular studies.

We can begin to build an integrated model to understand how *M. minutum* responds to two key stressors by integrating all the multi-omic information obtained, as outlined in Fig. 13. While we found that a number of genes contained common motifs (such as RTG, a stress factor that has been extensively studied in yeast [114] [115] [116]), we focused on the genes with the two most prominent motifs from our motif analysis ARF1 (highly enriched in scale-up stress) and an unannotated *Thalassiosira pseudonana* factor (T.p). We performed pathway analysis with the use of GPT-4, as suggested on [39], and see pathways consistent with the hubs of protein-protein interaction identified in Fig. 12. Taken together we have candidate genes, proteins and metabolites to target both for biomarker-based detection and for countermeasure and mitigation development.

**Fig. 12:**
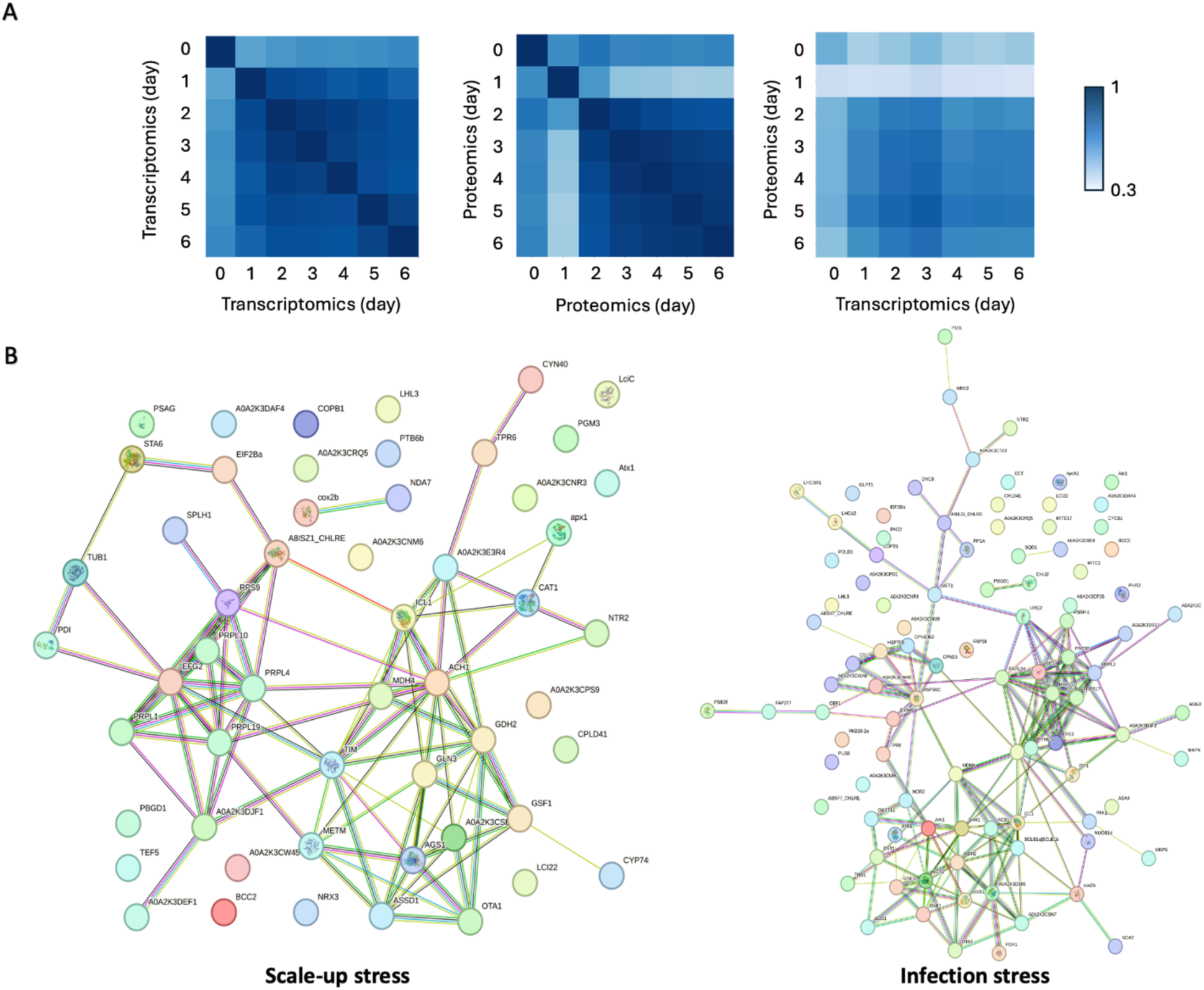
Understanding the dynamics between proteomic and transcriptomics observation and elucidating the individual network of features specifically related to the scale-up stress and the infection stress. A) Heatmap with the correlation between the proteomic and transcriptomic features. B) Network of protein features, significant for the scale-up and infection stress, made through STRING.

**Fig. 13:**
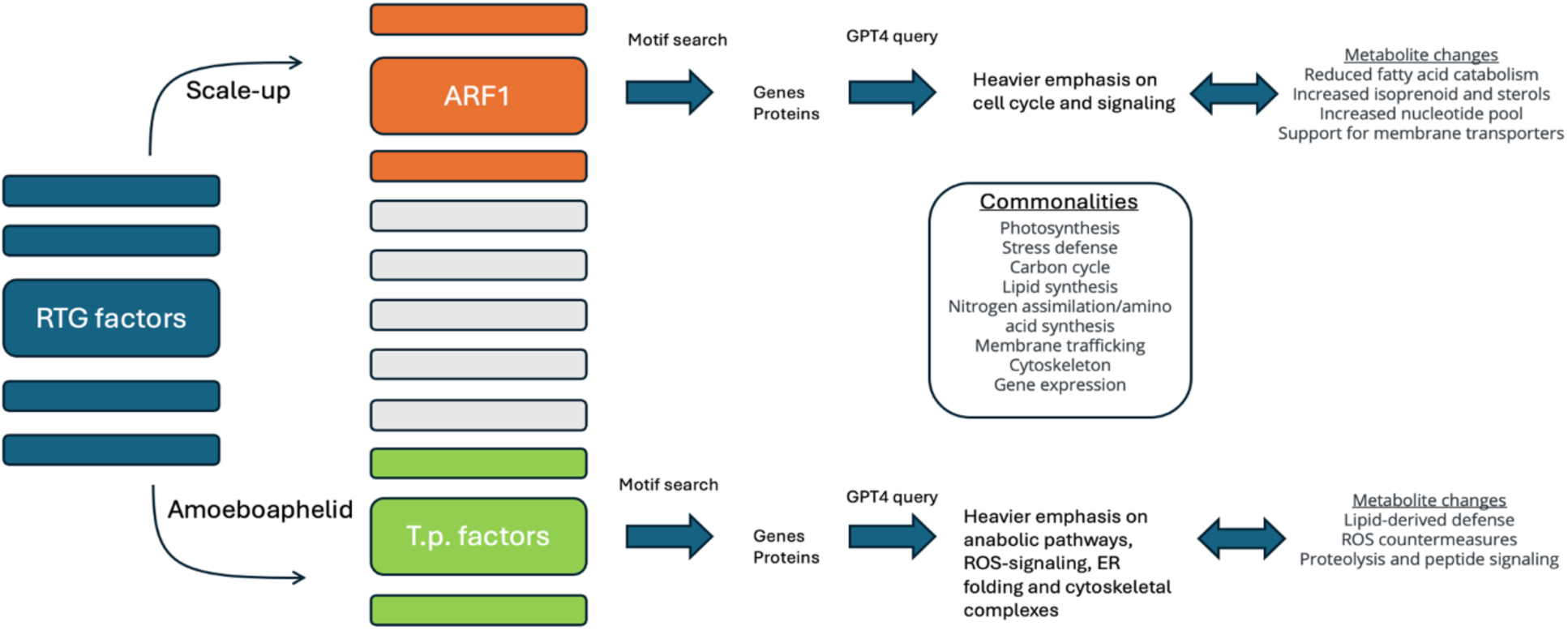
Procedural explanation of the use of GPT4 for stress-related regulation and metabolic changes.

With respect to metabolites in particular, for example, which are the most likely candidates for biomarker detection and immediate therapeutic intervention, specific hormones and steroids, such as 5-Dihydrotestosterone, 7-Hydroxytestosterone, Estrone, Estriol and 19-Norandrostenedion, are highly expressed during early infection. [73, 117, 118]Given that their abundance increased after the introduction of the pathogen, showcases that those compounds can be utilized as timely biomarkers for upcoming pond crashes. In the long term, if some of these molecules act in defense against algae, microbiomes can be engineered to produce these molecules as a prophylactic strategy during large-scale culture.

On a broader scale, the collection of molecular data from such expansive cultivation systems can pave the way for analyses aimed at developing predictive power with respect to culture resiliency, as well as potential targets for engineering this resiliency in the future. Indeed, we observed transcripts shared in both this study and the study conducted by Calhoun et al (2022), which indicates that more data generation in small- and large-scale systems, coupled with integrated analysis, could reveal novel health monitoring molecules in algal culture. The ultimate goal is for these models to be adaptable to multiple types of bioprocessing, beyond algae cultures, and this study provides a roadmap in that direction, using omic data to optimize large-scale applications.

## Abbreviations

14-HDHA: 14-hydroxy-docosahexaenoic acid
AFDW: Ash-free dry weight
ARF1: ADP-ribosylation factor 1
ARR1: Arabidopsis response regulator 1
CCM: Carbon concentration mechanism
DIABLO: Data Integration Analysis for Biomarker discovery using Latent Components
DISCOVR: Development of integrated screening, cultivar optimization, and verification research
DPGP: Dirichlet Process Gaussian Process mixture model
Msn4: multicopy suppressor of SNF1 mutation protein 4
PCA: principal component analysis
ROS: reactive oxygen species
RuBisCO: Ribulose-1,5-bisphosphate carboxylase/oxygenase
RTG1: mitochondrial retrograde 1
RTG3: mitochondrial retrograde 3
SNF1: sucrose non-fermenting 1

## Declarations

## Ethics approval and consent to participate

N/A

## Consent for publication

N/A

## Availability of data and materials

The datasets supporting the conclusions of this article are available in the NCBI SRA repository, under BioProject PRJNA1307767 and are included within the article and its additional files.

## Authors’ contributions

R.K. led the funding acquisition efforts and had the logistic and programmatic lead of the project. G.K and R.K. conceptualized the project and designed the methodology and the experimental procedure. G.K., D.Y., E.W., T.E., O.W. and M.P.H. worked on the different aspects of the experimental procedure (scale-up, pond running and maintenance, sampling and phenotyping, omics preparation and sequencing). G.K., J.S., D.Y., T.S. and R.K. worked on the computational and bioinformatic analysis. T.W.L. advised on the experimental design and algal physiology. G.K, J.S. and R.K. wrote the original manuscript. All the authors reviewed and edited the manuscript.

## Acknowledgements

This paper describes objective technical results and analysis. Any subjective views or opinions that might be expressed in the paper do not necessarily represent the views of the U.S. Department of Energy or the United States Government. We thank Ryan W. Davis, Catherine Mageeney, Pamela Lane, and Carolyn Fisher, Sharon Nademanee, and Damian Carrieri for their insightful comments on an earlier draft of this manuscript.

## Additional data

**Additional Figure. 1:**
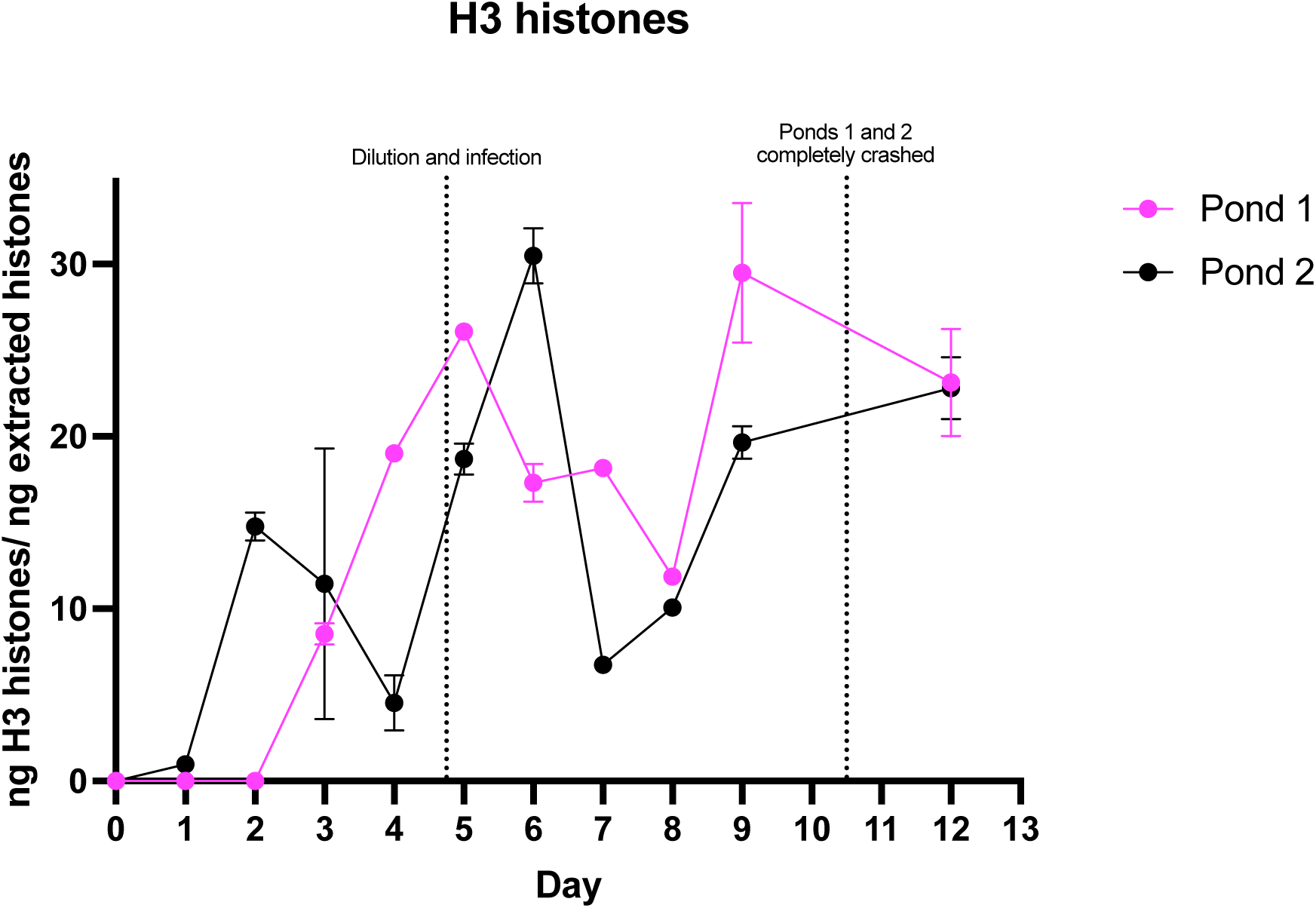
The caspase activity during the pond run. The caspase activity for every day of the culture run in pond 1 (pink) and pond 2 (black).

**Additional Figure 2:**
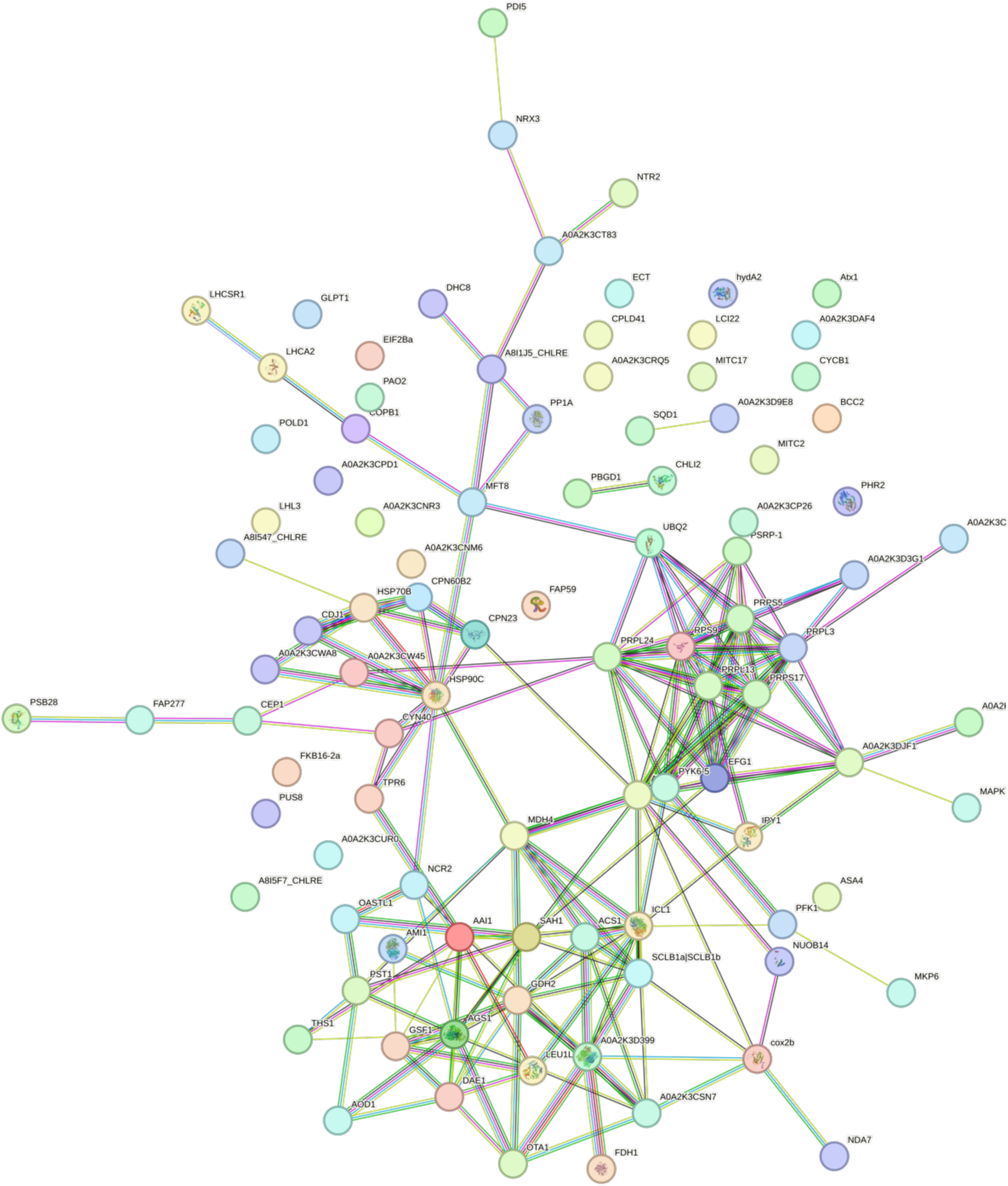
The complete STRING protein-protein network for the significant features during the scale-up stress.

**Additional Figure 3:**
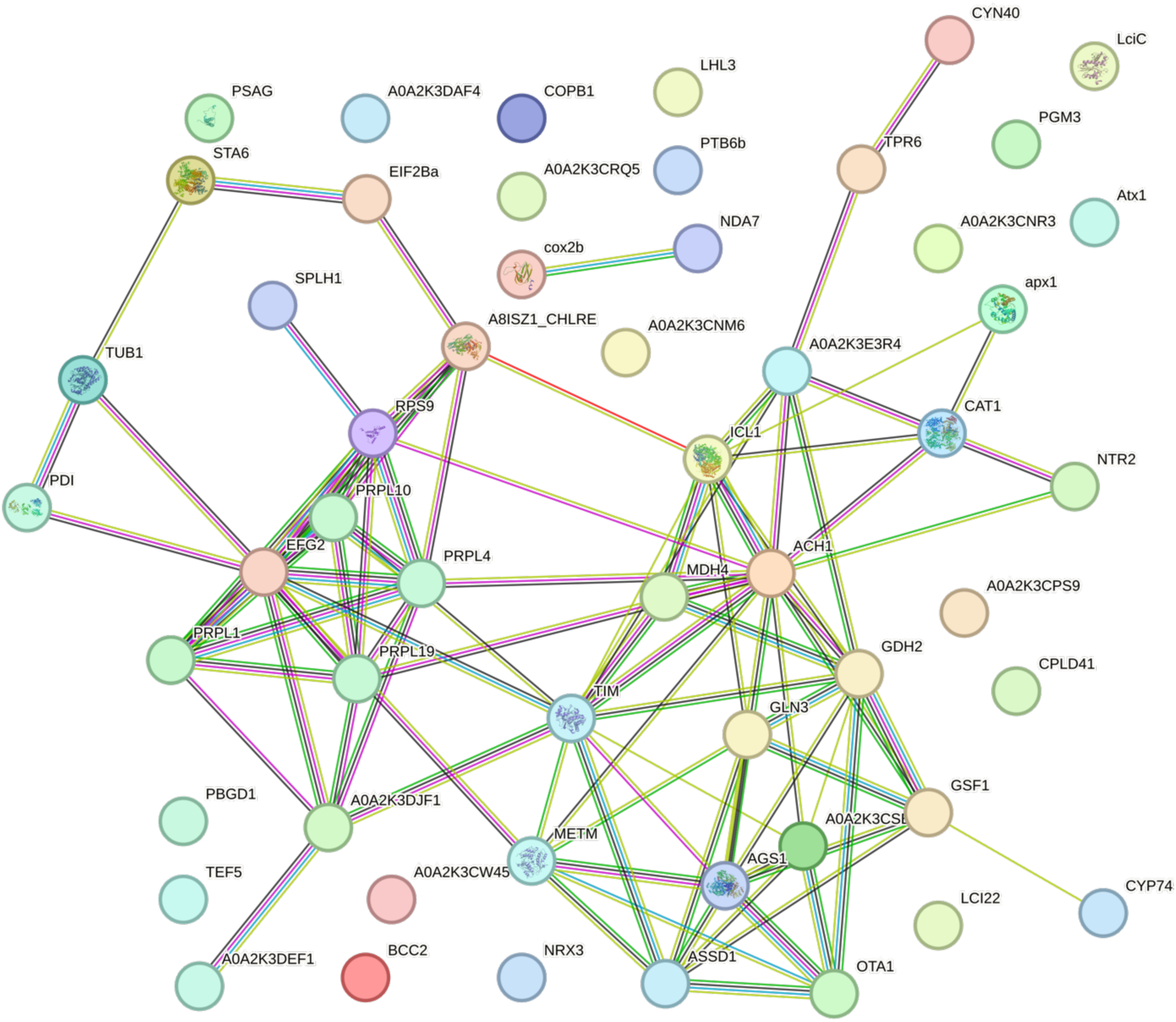
The complete STRING protein-protein network for the significant features during the infection stress.

Additional Dataset 1: Phenomics

Additional Dataset 2: Metagenomics

Additional Dataset 3: Transcriptomics

Additional Dataset 4: Proteomics

Additional Dataset 5: Metabolomics

Additional Dataset 6: Metabolomics small scale

